# Random Compressed Coding with Neurons

**DOI:** 10.1101/2022.01.06.475186

**Authors:** Simone Blanco Malerba, Mirko Pieropan, Yoram Burak, Rava Azeredo da Silveira

## Abstract

Classical models of efficient coding in neurons assume simple mean responses—‘tuning curves’—such as bell-shaped or monotonic functions of a stimulus feature. Real neurons, however, can be more complex: grid cells, for example, exhibit periodic responses which impart the neural population code with high accuracy. But do highly accurate codes require fine tuning of the response properties? We address this question with the use of a benchmark model: a neural network with random synaptic weights which result in output cells with irregular tuning curves. Irregularity enhances the local resolution of the code but gives rise to catastrophic, global errors. For optimal smoothness of the tuning curves, when local and global errors balance out, the neural network compresses information from a high-dimensional representation to a low-dimensional one, and the resulting distributed code achieves exponential accuracy. An analysis of recordings from monkey motor cortex points to such ‘compressed efficient coding’. Efficient codes do not require a finely tuned design—they emerge robustly from irregularity or randomness.

## 1 Introduction

Neurons convey information about the physical world by modulating their responses as a function of parameters of sensory stimuli. Classically, the mean neural response to a stimulus—referred to as the neuron’s ‘tuning curve’—is often described as a smooth function of a stimulus parameter with a simple monotonic or unimodal form (Georgopoulos et al., 1982; Taube et al., 1990; Miller et al., 1991; Bremmer et al., 1997; Dayan & Abbott, 2001; Kayaert et al., 2005). The deviation from the mean response—the ‘neural noise’—may lead to ambiguity in the identity or strength of the encoded stimulus, and the coding performance of a population of neurons as a whole is dictated by the forms of the tuning curves and the joint neural noise. In the study of population codes, the efficient coding hypothesis has served as a theoretical organizing principle. It posits that tuning curves are arranged in such a way as to achieve the most accurate coding possible given a constraint on the neural resources engaged (Barlow, 1961; Atick & Redlich, 1990; Lewicki, 2002). The latter is often interpreted as a metabolic constraint on the maximum firing rate of a single neuron or on the mean firing rate of the whole population (Zhang & Sejnowski, 1999; Bethge et al., 2002; Wang et al., 2016).

In order to tackle this constrained optimization problem in practice, tuning curves are parametrized, and the corresponding parameters are optimized. Here, the simplicity of the form of tuning curves matters: only a few parameters need to be optimized. A large body of literature addresses this constrained optimization problem, in particular in the perceptual domain. For example, many studies model tuning curves as Gaussian or other bell-shaped functions, and obtain the values of their centers and widths that minimize the ‘perceptual’ error committed when information is decoded from the activity of a population of model neurons (Zhang & Sejnowski, 1999; Deneve et al., 1999; Yaeli & Meir, 2010; Ganguli & Simoncelli, 2014; Fiscella et al., 2015). In the resulting optimal populations, and if noise among neurons is independent, the coding error typically scales like 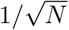, where *N* is the number of model neurons (Seung & Sompolinsky, 1993). This behavior can be intuited based on the observation that the ‘signal’ in the neural population grows like N while the noise grows like 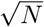, yielding a signal-to-noise ratio that increases in proportion to the square root of the population size. (In some models of population neural coding of a one-dimensional parameter, the width of tuning curves can be further optimized to yield an additional factor of 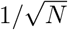; the error then scales like 1/*N* (Berens et al., 2011; Kim et al., 2020).)

Real neurons, however, can come with much more complex tuning curves than simple Gaussian or bell-shaped ones. Grid cells recorded in the enthorinal cortex offer a salient example (Hafting et al., 2005; Doeller et al., 2010; Yartsev et al., 2011; Killian et al., 2012); their tuning curves in two-dimensional, open field environments, are multimodal and periodic as a function of spatial coordinates. It was noted early on that such richer tuning curves can give rise to greatly enhanced codes. Given the periodicity of their tuning curves, and provided that the neural population includes several modules made up of cells with different periodicities (Fiete et al., 2008; Wei et al., 2015), grid cells can represent spatial location with an accuracy that scales exponentially (rather than algebraically, as above) in the number of neurons (Sreenivasan & Fiete, 2011; Mathis et al., 2012; Burak, 2014). Thus, the richer structure of individual tuning curves can be traded for a strong boost in the efficiency of the population code. Recent observations showed that place cells can also exhibit complex tuning curves in the context of motion in three dimensions, with multiple place fields that are irregular both in location and in size (Eliav et al., 2021). In addition, Ginosar et al. (2021); Grieves et al. (2021) found that during motion in three dimensional space, individual grid cells also exhibit irregular firing fields. A number of other examples of neurons with complex, but unstructured, tuning curves has also been identified in other cortical regions and in different species (Kadia & Wang, 2003; Sofroniew et al., 2015; Lalazar et al., 2016; Gaucher et al., 2020).

Here, we ask whether highly efficient codes must rely on finely-tuned properties, such as the tuning curves’ periodicity or the arrangement of different modules in the population, or, alternatively, arise generically and robustly in populations of neurons with complex tuning curves, in the absence of any fine tuning. We approach the question by studying the benchmark case of a random neural code: a population code which relies on irregular tuning curves that emerge from a simple, feedforward, shallow network with random synaptic weights. The input layer in the network is made up of a large array of ‘sensory’ neurons with classical, bell-shaped tuning curves; these neurons project onto a small array of ‘representation’ neurons with complex tuning curves. We show that, in the resulting population code, the coding error is suppressed exponentially with the number of neurons in this population, even in the presence of high-variance noise.

In the context of this highly efficient code, it is not sufficient to consider a ‘typical error’: efficiency results from the compression of the stimulus space into the activity of a layer of neurons of comparatively small size; the price to pay for this compression is the emergence of two qualitatively distinct types of error—‘local errors’, in which the encoding of nearby stimuli is ambiguous, and ‘global (or catastrophic) errors’, in which the identity of the stimulus is lost altogether. The efficient coding problem then translates into a trade-off between these two types of errors. In turn, this trade-off yields an optimal width of the tuning curves in the ‘sensory layer’: when stimulus information is compressed into a ‘representation layer’, tuning curves in the sensory layer have to be sufficiently wide as to prevent a prohibitive rate of global errors.

We first develop the theory for a one-dimensional input (e.g., a spatial location along a line or an angle), then generalize it to higher-dimensional inputs. The latter case is more subtle because the sensory layer itself can be arranged in a number of ways (while still operating with simple, classical tuning curves). This generalization allows us to apply our model to data from monkey motor cortex, where cells display complex tuning curves. We fit our model to the data and discuss the merit of a complex ‘representation code’. Overall, our approach can be viewed as an application of the efficient coding principle to a framework that includes a downstream (‘representation’) layer of neurons as well as a peripheral (‘sensory’) layer of neurons. Our study extends earlier theoretical work on grid cells and other ‘finely designed’ codes by proposing that efficient compression of information can occur robustly even in the case of a random network. We reach our results by considering the geometry of population activity in a compressed, representation layer of neurons.

## 2 Results

We organize the description of our results as follows. First, we present, in geometric terms, the qualitative difference between a code that uses simple, bell-shaped tuning curves and one that uses more complex forms. Second, we introduce a simple model of a shallow, feedforward network of neurons that can interpolate between simple and complex tuning curves depending on the values of its parameters. Third, we characterize the accuracy of the neural code in the limiting case of maximally irregular tuning curves. Fourth, we extend the discussion to the more general case in which an optimal code is obtained from a trade-off between local and global errors. All the above is done for the case of a one-dimensional input space. Fifth, we generalize our approach to the case of a multi-dimensional stimulus. This allows us, sixth, to apply our model to recordings of motor neurons in monkey, and to analyze the nature of population coding in that system. Seventh, we give a quantitative description of the geometry of the population response induced by our network as a function of its parameters, through a measure of dimensionality. Finally, we extend our model to include an additional source of noise—‘input noise’ in the sensory layer, in addition to the ‘output noise’ present in the representation layer; input noise gives rise to correlated noise downstream, and we analyze its impact on the population code.

### The geometry of neural coding with simple vs. complex tuning curves

A neural code is a mapping that associates given stimuli to a probability distribution on neural population activity; in particular, the code maps any given stimulus to a mean population activity. In the case of a continuous, one-dimensional stimulus space, the latter is mapped into a curve in the *N*-dimensional space of the population activity, whose shape is dictated by the form of the tuning curves of individual neurons. As an illustration, we compare the cases of three neurons with Gaussian tuning curves and three neurons with periodic (grid-cell-like) tuning curves with three different periods (Fig. 1A). Simple tuning curves generate a smooth population response curve, implying that similar stimuli are mapped to nearby responses; by contrast, more complex tuning curves give rise to a serpentine curve. The latter makes better use of the space of possible population responses than the former, and hence can be expected to yield higher-resolution coding. Indeed, when the population response is corrupted by noise of a given magnitude, it will elicit a smaller *local* error in the case of complex tuning than in the case of simple tuning: by ‘stretching’ the mean response curve over a longer trajectory within the space of possible population activities, complex tuning affords the code with higher resolution relative to the range of the encoded variable. However, this argument does not capture in full the influence of noise on the nature of coding errors. In the case of a winding and twisting mean response curve, two distant stimuli are sometimes mapped to nearby activity patterns. In the presence of noise, this geometry gives rise to *global* (or catastrophic) errors. The enhanced resolution of the neural code associated with the occurrence of global errors was also noted in the context of grid-cell coding (Welinder et al., 2008; Sreenivasan & Fiete, 2011). Because of this trade-off, whether a simple or complex coding scheme is preferable becomes a quantitative question, which depends upon the details of the structure of the encoding.

**Figure 1:**
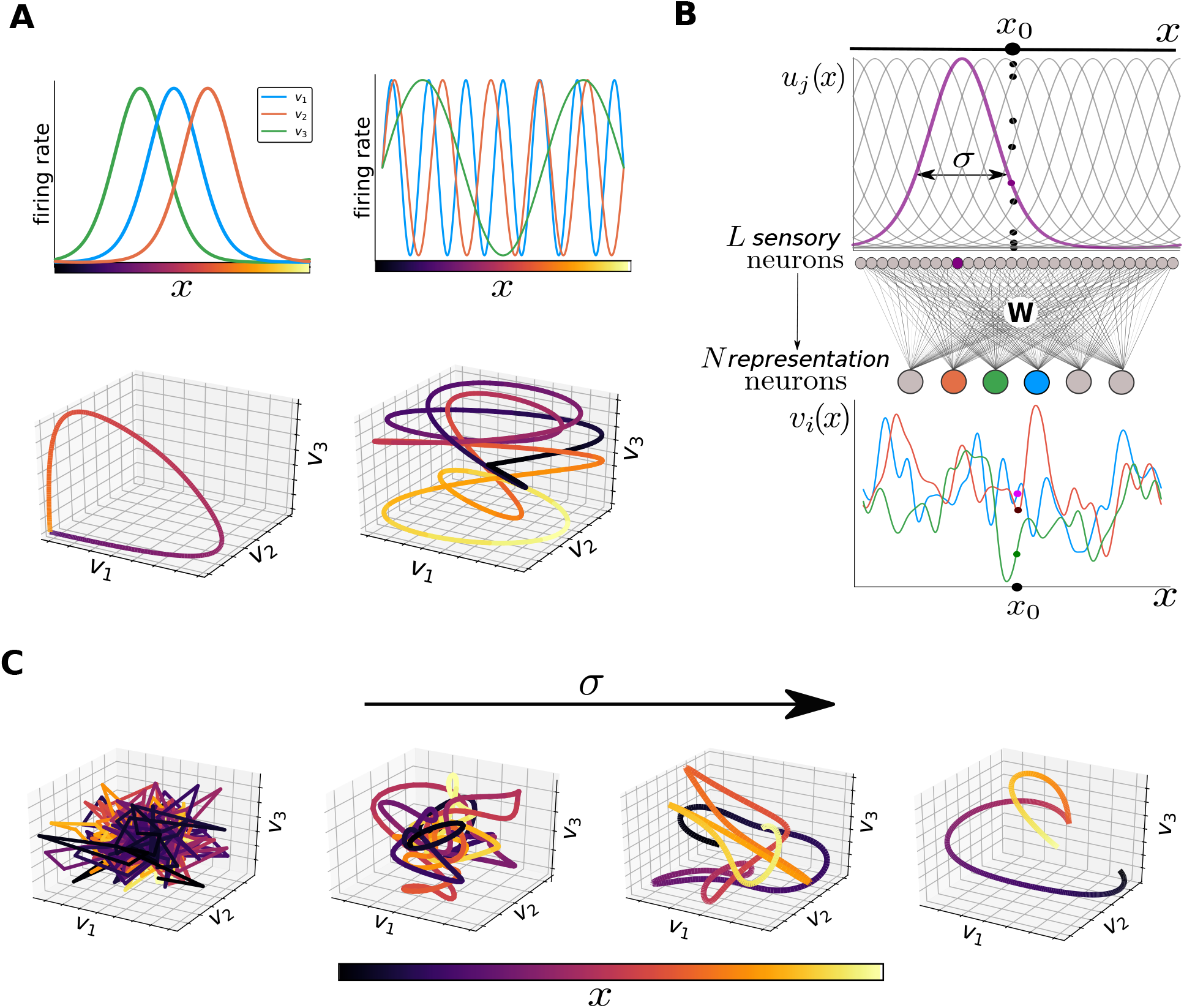
Geometrical approach to coding, and the random feedforward neural network architecture. (**A**) Top: mean responses of three-neuron populations encoding a one-dimensional stimulus. Left: population of neurons with Gaussian tuning curves. Right: population of neurons with periodic tuning curves. Bottom: mean activity in the neural populations, parametrized by the stimulus value colored according to the legend, as one-dimensional curves in a three-dimensional space. Unimodal tuning curves (left) evoke a single-loop curve, which preserves the distances between stimuli in the evoked responses. Periodic tuning curves (right) evoke a more complex curve in which two distant stimuli may be mapped to nearby points in the joint-activity space; the curve is longer, and fills up a larger portion of the activity space. (**B**) Feedforward neural network. An array of *L* sensory neurons with Gaussian tuning curves (one highlighted in purple) encodes a one-dimensional stimulus into a L-dimensional representation. These tuning curves determine the mean response of the population for a given stimulus, *x_0_* (dots). This layer projects onto a smaller layer of *N* representation neurons with an all-to-all random-connectivity matrix, **W**, generating irregular responses. We plot the tuning curves of three sample neurons, highlighting their response to the stimulus *x*_0_. (**C**) Examples of population activity (across the stimulus line, color indicates stimulus value) for three sample representation neurons, for increasing values of *σ*. When *σ* → 0 (left, *σ* = 0.001), neurons produce uncorrelated random responses to different stimuli, generating a spiky curve made up by broken segments. As *σ* grows (*σ* = 0.015, *σ* = 0.03) irregularities are smoothed out, and nearby stimuli evoke increasingly correlated responses. Ultimately, for large values of *σ* (right, *σ* = 0.15) we recover a scenario similar to that with unimodal tuning curves.

### Shallow feedforward neural network as a benchmark for efficient coding

In order to address the problem mathematically, we examine the simplest possible model that generates complex tuning curves, namely a two-layer feedforward model. An important aspect of the model is that it does not rely on any finely-tuned architecture or parameter tuning: complex tuning curves emerge solely because of the variability in synaptic weights; thus, the model can be thought of as a benchmark for the analysis of population coding in the presence of complex tuning curves. The architecture of the model network and the symbols associated with its various parts are illustrated in Fig. 1B. In the first layer, a large population of *L sensory* neurons encodes a one-dimensional stimulus, *x*, into a high-dimensional representation. Throughout, we assume that *x* takes values between zero and one, without loss of generality. (If the input covered an arbitrary range, say *r*, then the coding error would be expressed in proportion to *r*. In other words, one cannot talk independently of the range of the input and of the resolution of the code. We set the range to unity in order to avoid any ambiguity.) Sensory neurons come with classical tuning curves: the mean activity of neuron *j* in response to stimulus *x* is given by a Gaussian with center *C_j_* (the preferred stimulus of that neurons) and width *σ*:

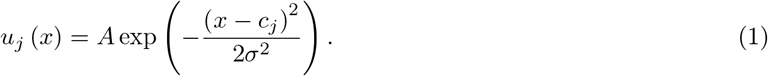

Following a long line of models, we assume that the preferred stimuli in the population are evenly spaced, so that *C_j_* = *j/L*. As a result, the response vector for a stimulus *x*_0_, **u** (*x*_0_), can be represented as a Gaussian ‘bump’ of activity centered at *x*_0_.

Complex tuning curves appear in the second layer containing *N representation* neurons; we shall be interested in instances with *N* ≪ *L*, in which efficient coding results in compression of the stimulus information from a high-dimensional to a low-dimensional representation. Each representation neuron receives random synapses from each of the sensory neurons; specifically, the elements of the all-to-all synaptic matrix, **W**, are i.i.d. Gaussian random weights with vanishing mean and variance equal to 1/*L* (*W_ij_* ~ *N* (0,1/*L*)). In the simple, linear case that we consider, the mean activity of neuron *i* in the second layer is thus given by

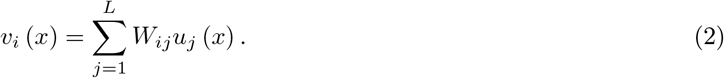

Since the weights *W_ij_* correspond to a given realization of a random process, they generate tuning curves, *v_i_* (*x*), with irregular profiles. The parameter *σ* is important in that it controls the smoothness of the tuning curves in the second layer: it defines the width of *u_j_*, which in turn dictates the correlation between the values of the tuning curve *v_i_* for two different stimuli. By the same token, the amplitude of the variations of *v_i_* with *x* depends upon the value of *σ*. For a legitimate comparison of population codes in different networks, we set this amplitude to a constant on average,

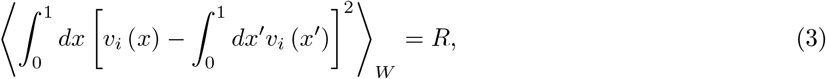

by calibrating the value of the prefactor in Eq. (1), *A*. Because of the averaging over the synaptic weights, indicated by the brackets 〈·〉_*W*_, A does not depend upon a specific realization of the synaptic weights. Equation (3) corresponds to the usual constraint of ‘resource limitation’ in efficient coding models; it amounts to setting a maximum to the variance of the output over the stimulus space, as is commonly assumed in analyses of efficient coding in sensory systems (Atick & Redlich, 1990; Van Hateren & Ruderman, 1998; Doi et al., 2012; Zhaoping, 2014).

Returning to our geometric picture, we observe that, by changing the value of *σ*, we can interpolate between smooth and irregular tuning curves in the second layer (Fig. 1C). In the limiting case of large *σ*, representation neurons come with smooth tuning curves akin to classical ones; in the other limiting case of small *σ*, the mean population response curve becomes infinitely tangled. Thus, as the value of *σ* is decreased, the mean response curve ‘stretches out’ and necessarily twists and turns, in such a way as to fit within the allowed space of population responses defined by Eq. (3). A longer population response curve fills the space of population responses more efficiently and represents the stimulus at a higher resolution, but its twists and turns may result in greater susceptibility to noise.

To complete the definition of the model, we specify the nature of the noise in the neural response. We assume that the activity of neuron *i* in the second layer is affected by noise, which we denote by *z_i_*, such that its response at each trial (in which stimulus *x* is presented) is given by *r_i_* = *v_i_* (x) + *z_i_*. For the sake of simplicity, we use Gaussian noise with vanishing mean and variance equal to *η*^2^. In most of our analyses, we suppose that responses in the first layer are noiseless and that the noise in the second layer is uncorrelated among neurons; in the last subsection, however, we relax these assumptions, and discuss the implications of noisy sensory neurons and correlated noise among representation neurons. (Our motivation for considering noiseless sensory neurons is that we are primarily interested in analyzing the compression of the representation of information between the first and the second layer of neurons. By contrast, noise in sensory neurons affects the fidelity of encoding in the *first* layer already.)

We quantify the performance of the code in the second layer through the mean squared error (MSE) in the stimulus estimate as obtained from an ideal decoder, ‘ideal’ in the sense that it minimizes the MSE. (Throughout, in heuristic arguments and analytical calculations, we focus on the MSE. In a number figures, however, we plot its square root, the RMSE, so as to allow for a direct comparison with the stimulus range. The figure captions specify which of the two quantities is illustrated.) The use of an ideal decoder is an abstract device that allows us to focus on the uncertainty inherent to *encoding* (rather than to imperfections in *decoding*); it is nevertheless possible to obtain a close approximation to an ideal decoder in a simple neural network with biologically plausible operations (see Methods).

### Compressed coding in the limiting case of narrow sensory tuning

It is instructive to study the properties of coding in our model in the limiting case of neurons with narrow tuning curves in the sensory layer (*σ* → 0), because this limit yields the most irregular tuning curves in the representation layer of our network (Fig. 1C). As we shall see, this limiting case also corresponds to that of a completely uncorrelated, random code, for which the mathematical analysis simplifies. When the value of *σ* is much smaller than 1/*L*, neurons in the sensory layers respond only if the stimulus coincides with the preferred stimulus of one of the neurons, and only that neuron is activated by the stimulus presentation; stimulus values that lie in between the preferred stimuli of successive sensory neurons in the first layer do not elicit any activity in the system. We can thus consider that any stimulus of interest is effectively chosen within a discrete set of L stimuli with values *x_j_* = *j/L*, with *j* = 1,..., *L*.

Each of these stimuli elicits a mean response

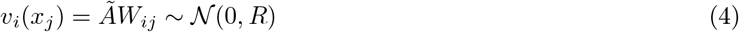

in neuron *i* of the second layer. Here, the value of *Ã* is chosen so as to set the amplitude of the variations of *v_i_* to be equal to the constant *R* (analogously to Eq. (3) but for the case of discrete stimuli). Geometrically, Eq. (4) represents a mapping from L stimulus values to a set of uncorrelated, random locations in the space of the population activity (as illustrated in Fig. 2A for a two-neuron population). In any given trial, however, the responses in the representation layer are corrupted by noise (Fig. 2A). The ideal decoder interprets a single-trial response as being elicited by the stimulus associated to the nearest possible mean response (Fig. 2A). The outcome of this procedure can be twofold: either the correct or an incorrect stimulus is decoded; in the latter case, because the possible mean responses are arranged randomly in the space of population activity (Fig. 2A and Eq. (4)), errors of all magnitudes are equiprobable. In other words, a model with narrow sensory tuning curves results in a second-layer code that does not preserve distances among inputs, and, consequently, the decoding error is either vanishing or, typically, on the order of the input range (set to unity here). The mean error is then simply proportional to the probability with which the ideal decoder makes a mistake, with a constant of proportionality of the order of the stimulus range.

**Figure 2:**
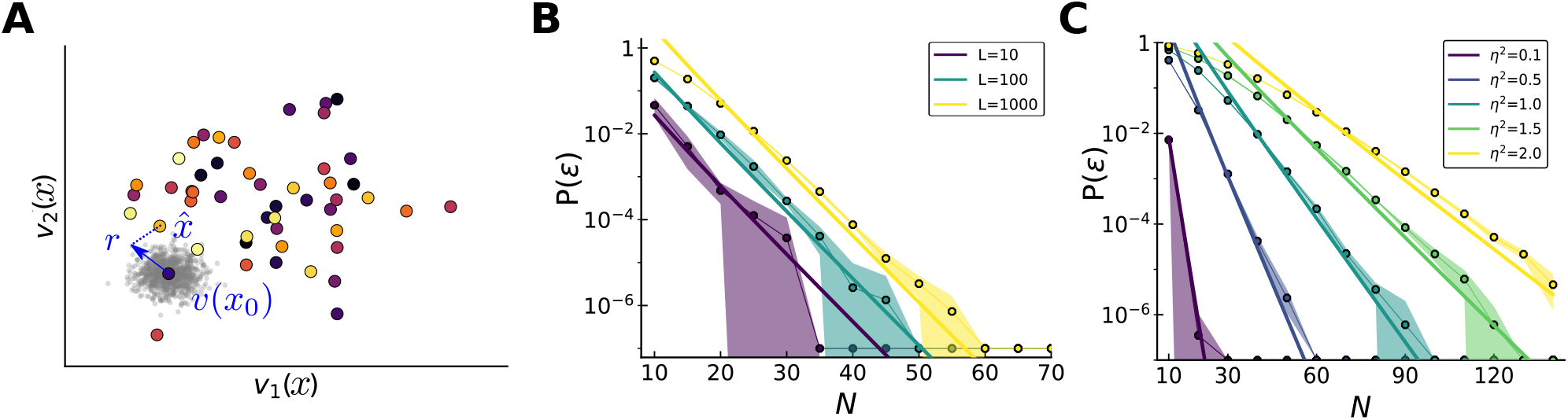
Probability of error for narrow tuning curves in the sensory layer. (**A**) Joint mean responses of two neurons to *L* = 50 stimuli, colored according to the legend in Fig. 1C. Noise is represented as a cloud of possible responses (in grey) around the mean. An error occurs when the noisy response, **r**, falls closer to a mean response corresponding to a stimulus, 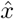, different from the true one, *x*_0_. Since mean responses are uncorrelated, 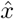 may be distant from *x*_0_. (**B**) Theoretical (solid curves, Eq. (5)) and numerical (dots) results for the probability of error as a function of the population size, for different values of *L* (*η*^2^ = 0.5). The probability of error scales exponentially with the number of neurons, *N*, with a multiplicative constant involving the number of stimuli, *L*. (**C**) Theoretical (solid curves) and numerical (dots) results for the probability of error as a function of the population size for different values of *η*^2^ (*L* = 500).

In Methods, we provide a derivation of this quantity. In the case of low-error coding, which interests us, we obtain the dependence of the probability of a decoding error as a function of the various model parameters, as

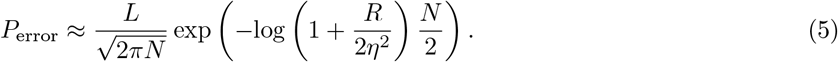

The main dependence to note, here, is the exponentially strong suppression as a function of the number of neurons in the second layer (Fig. 2B). By contrast, the probability of error scales merely linearly with the size of the stimulus space, *L*, as is expected in the low-error limit. This result implies that it is possible to compress information highly efficiently in a comparatively small representation layer (*N ≪ L*) *even though* the code is completely random. The price to pay for the use of randomness is that any error is likely ‘catastrophic’ (on the order of the stimulus range), but these large errors happen prohibitively rarely. It is also worth noting that the rate of exponential suppression depends on the variance of the noise, *η*^2^, or, more precisely, on the single-neuron signal-to-noise ratio, *R/η*^2^ (where *R* is the variance of the signal, Eq. (3)). In numerical simulations, we set *R* =1 and we vary *η*^2^ to explore different noise regimes. Interestingly, even when this signal-to-noise ratio becomes small, i.e., when the noise in the activity of individual neurons is comparable to modulations of their mean responses, the exponential suppression as a function of *N* of the probability of error remains valid, with a rate approximately equal to *R*/4*η*^2^.

### Compressed coding with broad tuning curves: trade-off between local and global errors

As we saw in the previous section, in the case of infinitely narrow tuning curves the coding of a stimulus in a given trial is either perfect or indeterminate; that is, any error is typically a global error, on the order of the entire stimulus range. In the more general case of sensory neurons with arbitrary tuning width, the picture is more complicated: in addition to *global* errors which result from the twisting and turning of the mean response curve, the population code is also susceptible to *local* errors (Fig. 3A). This is because broad tuning curves in the sensory layer partly preserve distances: locally, nearby stimuli are associated with nearby points on the mean response curve; as a result, the coding of any given stimulus is susceptible to local errors due to the response noise. As the tuning width in the sensory layer, *σ*, decreases, two changes occur in the mean response curve: it becomes longer (it ‘stretches out’) and it becomes more windy (Fig. 1C). Stretching increases the local resolution of the code (because it allows for two nearby stimuli to be mapped to two more distant points in the space of population activity), while windiness increases the probability of global errors. This trade-off is apparent when we plot the histogram of error magnitudes as a function of *σ*: for larger values of *σ*, global errors are less frequent, but local errors are boosted (Fig. 3B). Also noticeable, here, is that the large-error tails of the histograms are flat, consistent with the observation that global errors of all sizes are equiprobable. (Strictly speaking, this happens if the stimulus has periodic boundary conditions, such that, picking two random points, the probability that they are at a given distance does not depend on the location of one or the other point.) For a more quantitative understanding, we carried out an approximate analytical calculation, in which *(i*) we approximated the mean response curve by a linear function locally and (ii) we considered that the distance between two segments of the curve representing the mean responses to two stimuli distant by more than *σ* is random and independent of the stimulus values. Using these two assumptions, we obtained the MSE as a sum of two terms (see Methods for mathematical details) corresponding to local and global errors, as

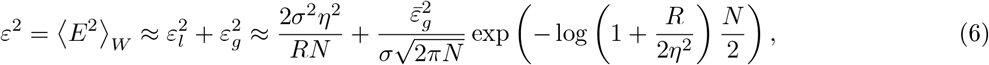

where 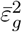 is a term of 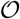 (1) that depends upon the choice of stimulus boundary conditions (see Methods). This expression quantifies the MSE for a ‘typical’ network, obtained by averaging over possible choices of synaptic weights, as indicated by the brackets 〈·〉_*W*_. The first term on the right-hand-side of Eq. (6) represents the contribution of local errors, while the second term corresponds to global errors (Fig. 3C). Their form can be intuited as follows. The magnitude of local errors is proportional to *η*^2^ and inversely proportional to *N*, as in classical models of population coding with neurons with bell-shaped tuning curves (see, e.g., Zhang & Sejnowski (1999)). Furthermore, decreasing *σ* stretches out the mean response curve, which increases the local resolution of the code and explains the factor *σ*^2^ in Eq. (6). (The form of this first term can also be understood as the inverse of the Fisher information (Seung & Sompolinsky, 1993; Brunel & Nadal, 1998), which bounds the variance of an unbiased stimulus estimator.) The second term on the right-hand-side of Eq. (6) is obtained as an extension of Eq. (5): instead of considering the probability that two mean response points are placed nearby, we consider the probability that two segments of the mean response curve with size *σ* each fall nearby. There are 1/*σ* such segments (since we have set the stimulus range to unity), and this explains why the factor *L* in Eq. (5) is replaced by a factor 1/*σ* in Eq. (6). Importantly, the two terms in Eq. (6) are modulated differently by the two parameters *N* and *σ*. Depending upon their values, either local or global errors dominate (Fig. 3C).

**Figure 3:**
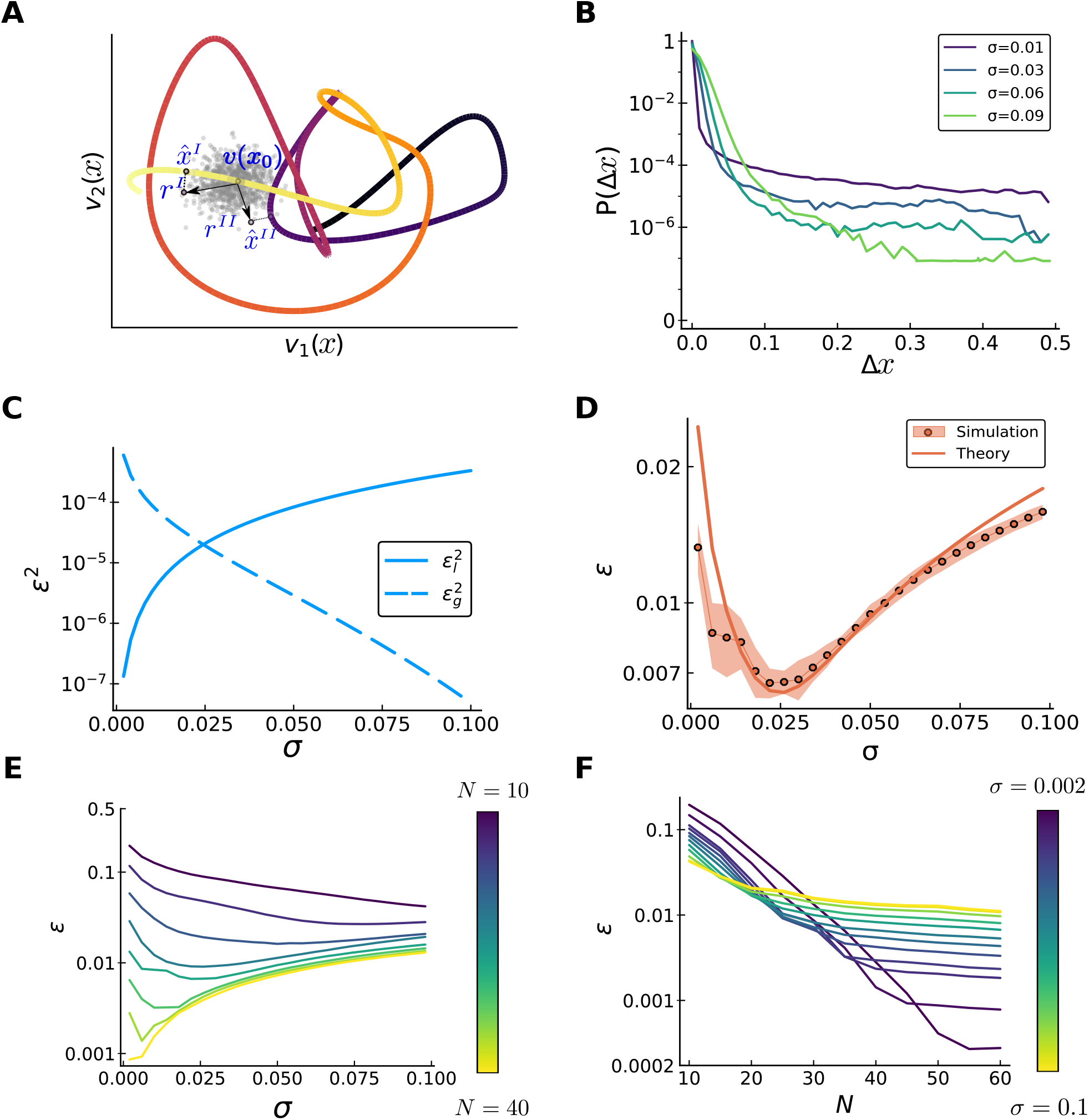
Trade-off between local and global errors. *continue to next page* Figure 3 *(previous page)*: (**A**) Different types of error in an irregular curve of mean population activity (joint response of two neurons, colored according to the legend in Fig. 1C). Here, **r**^*I*^ and **r**^*II*^ are two possible noisy responses to the same stimulus, extracted from the Gaussian cloud surrounding the mean response, **v**(*x*_0_). An ideal decoder outputs the stimulus corresponding to the closest point on the curve. In one case, **r**^*I*^ results in a local error, by selecting a point on the curve that represents a nearby stimulus, 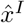. In the other case, **r**^II^ is closer to a point on the curve which represents a stimulus distant from the true one, 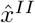, causing a global error. (**B**) Normalized histogram of absolute error magnitudes, 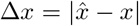, made by an ideal decoder, for different values of *σ* (*N* = 25). For better visualization, we consider a stimulus with periodic boundary conditions. The contribution of the two types of error varies with *σ*. For small *σ*, coding is precise locally (fast drop of the purple curve for small errors), but many global errors occur (tail of the distribution is high). For large *σ* (green curves) local accuracy is poorer but global errors are suppressed. (**C**) Theoretical prediction for the two contributions to the MSE as a function of *σ* (*N* = 30). The magnitude of local errors increases with larger *σ* (solid curve), while the number of global errors decreases (dashed curve). (**D**) RMSE as a function of *σ*: comparison between numerical simulations (dots) and theoretical prediction of Eq. (6) (solid curve). (**E**) RMSE, as a function of *σ* for different population sizes N (increasing from violet to yellow). The smallest RMSE occurs at an optimal value of *σ*, *σ*\*(*N*), which decreases with increasing *N*. (**F**) Same data, but the error is displayed as a function of *N*, for a fixed value of *σ*. The MSE decreases exponentially rapidly until global errors are suppressed, then the local errors are linearly reduced. A smaller value of *σ* implies a larger value of *N* at which the crossover occurs, as well as a smaller MSE at this crossover value.

We tested the validity of Eq. (6): it agrees closely with results from numerical simulations, in which we computed the MSE using a Monte Carlo method and a network implementation of the ideal decoder (Fig. 3D, see Methods for details). The non-trivial dependence is illustrated by the observation that the MSE may decrease or increase as a function of *σ*, around a given value of *σ*, depending upon the value of *N* (Fig. 3E). Furthermore, the strong (exponential) reduction in MSE with increasing *N* occurs only up to a crossover value that depends on *σ* (Fig. 3F); beyond this value, global errors disappear, and the error suppression is shallower (hyperbolic in *N*, due to improved local resolution). For small values of *σ*, the crossover values of *N* are larger and occur at lower values of the MSE.

As is apparent from Figs. 3D and E, for any value of *N* there exists a specific value of *σ* = *σ*\* (*N*) that balances the two contributions to the MSE such as to minimize it. This optimal width can be thought as the one that stretches out the mean response curve as much as possible to increase local accuracy but that stops short of inducing too many catastrophic errors. The MSE is asymmetric about the optimal width, *σ*\*: smaller values of *σ* cause a rapid increase of the error due to an increased probability of global errors, while larger values of *σ* mainly harm the code’s local accuracy, resulting in a milder effect. From Eq. (6), we obtain the dependence of the optimal width upon the population size, as

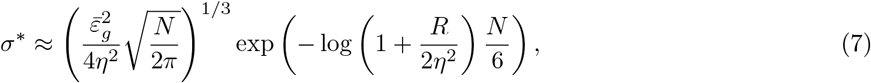

and the optimal MSE as a function of *N*, as

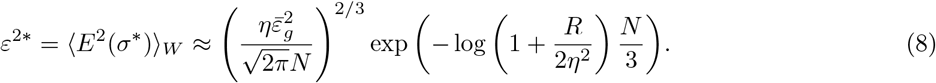

Both these analytical results agree closely with numerical simulations (Figs. 4A and B). Equation (8) and Fig. 4B show that the optimal MSE is suppressed exponentially with the number of representation neurons in the second layer. Thus, highly efficient compression of information and exponentially strong coding also occurs when tuning curves in the sensory layer are *not* infinitely narrow: furthermore, a degree of smoothness in the tuning of the sensory neurons is advantageous. With the optimal choice of the sensory tuning width, the rate of scaling with N of the argument within the exponential in Eq. (8) depends upon the noise variance, *η*^2^; in Figs. 4C and D, we illustrate the dependence of *σ*\* and *ε*\* upon *N* and *η*^2^.

**Figure 4:**
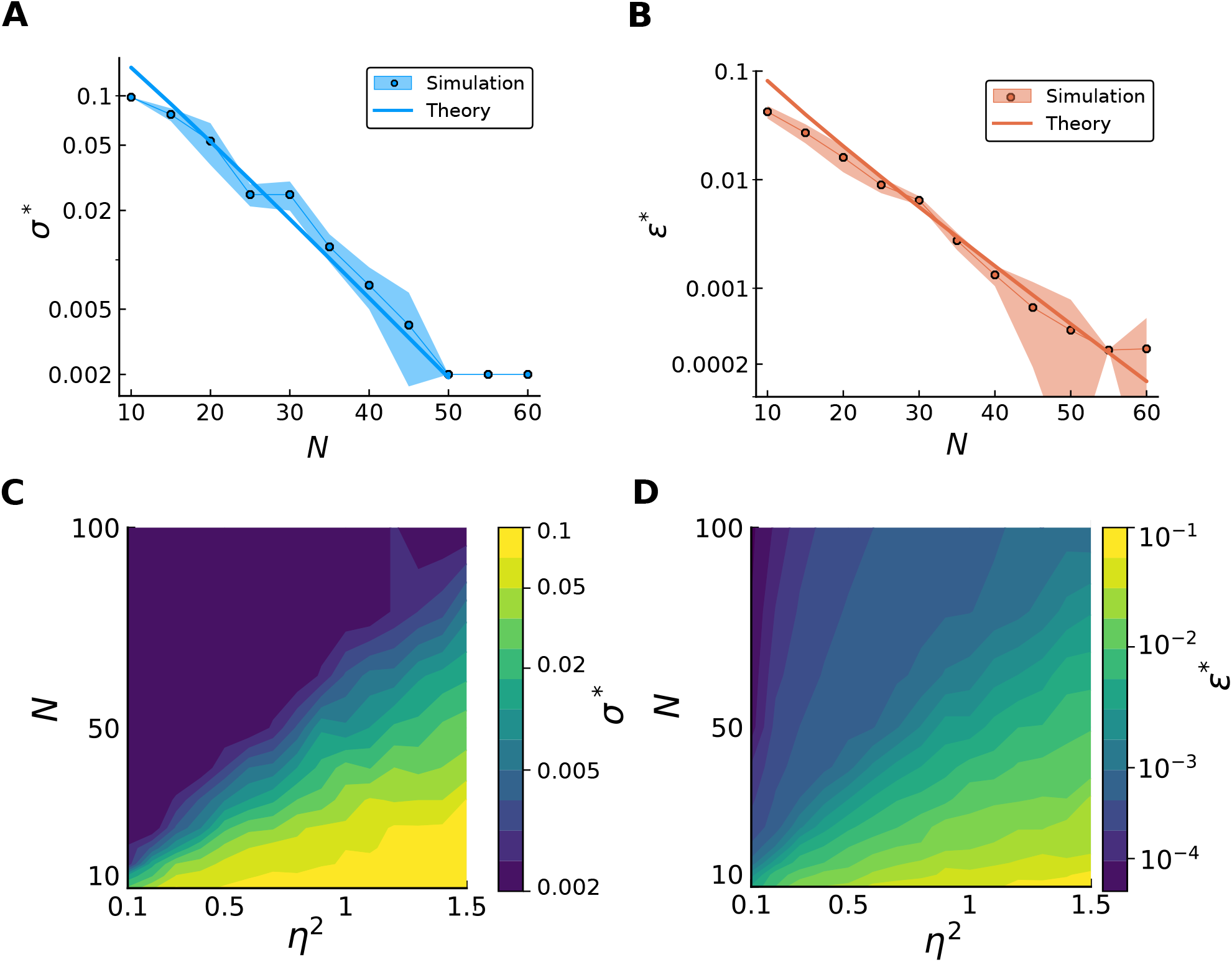
Scaling of the optimal width and the optimal MSE as a function of population size and signal-to-noise ratio. (**A**) The optimal *σ*\* decreases exponentially rapidly with the number of representation neurons, saturating the lower bound imposed by the finite number of neurons of the first layer (corresponding to the spacing of the preferred positions, 1/*L*). Simulations (dots) show good agreement with analytical results (solid curve). (**B**) The optimal RMSE is suppressed exponentially rapidly with *N*. Simulations (dots) agree with analytical results (solid curve). (**C,D**) Optimal width (**C**) and RMSE (**D**) as a function of the parameters *N* and *η*^2^. The color coding is in log scale, in order to highlight the exponential scaling.

### Compressed coding of multi-dimensional stimuli

Real-world stimuli are multi-dimensional. Our model can be extended to the case of stimuli of dimensions higher than one, but particular attention should be given to the nature of encoding in the first layer—because sensory neurons can be sensitive to one or several dimensions of the stimulus. In one limiting case, a sensory neuron is sensitive to all dimensions of the stimulus; for example, place cells respond as a function of the two- or three-dimensional spatial location. Visual cells constitute another example of multi-dimensional sensitivity, as they respond to several features of the visual world; for example, retinal direction-selective cells are sensitive to the direction of motion, but also to speed and contrast. In the other limiting case, sensory neurons are tuned to a single stimulus dimension, and insensitive to others. We will refer to these two coding schemes as *pure* and *conjunctive,* following Ref. Finkelstein et al. (2018) where they are examined in the context of head-direction neurons in bats. The authors conclude that the relative advantage of a pure coding scheme—with neurons that encode a single head-direction angle—with respect to a conjunctive coding scheme—with neurons that encode two head-direction angles—depends on specific contingencies, such as the population size or the decoding time window. Indeed, in a conjunctive coding scheme individual neurons carry more information, but the population as a whole needs to include sufficiently many neurons to cover the (multi-dimensional) stimulus space—a constraint which becomes more restrictive as the number of dimensions increases.

We generalized our model to include the possibility of *K*-dimensional stimuli. For the sake of simplicity, we consider here only the two limiting cases of *pure* and *conjunctive* coding in the *sensory* layer of our model (i.e., we do not discuss intermediate cases, in which a given sensory neuron is sensitive to several but not all stimulus dimensions, see Methods). In the model, furthermore, neurons in the *representation* layer receive random inputs from *all* sensory neurons; as such, the representation layer always embodies a conjunctive coding scheme.

By extending the geometric picture (illustrated in Fig. 1 for the case of a one-dimensional stimulus), we can analyze differences in coding properties between pure and conjunctive coding schemes; in Fig. 5A, we illustrate the case of a two-dimensional stimulus. In this case, the mean response of representation neurons corresponds to a mapping from a two-dimensional stimulus space to a random ‘sheet’ (a two-dimensional surface) in the N-dimensional space of the population activity. In the *pure case,* the activity of a given sensory neuron is maximally modulated when the stimulus varies along a particular dimension, the one to which the neuron is sensitive. Variations of the stimulus along orthogonal directions have no effect on the mean neural activity. Neurons in the representation layer compute a linear sum of these responses, and therefore their activity can be decomposed as a sum of one-dimensional functions. This implies that the ‘response sheet’ is maximally curved along each of the stimulus dimensions; geometrically, this results in a ‘folded’ structure, with creases along the directions of mild sensitivity. By contrast, in the *conjunctive case* the activity of a sensory neuron is modulated by variations of the stimulus along any direction. As a result, the ‘response sheet’ that represents the joint mean activity of neurons in the second layer comes with (random) curvature equally along all stimulus dimensions: rather than ‘folded’, it behaves like a ‘crumpled’ sheet (Fig. 5A).

**Figure 5:**
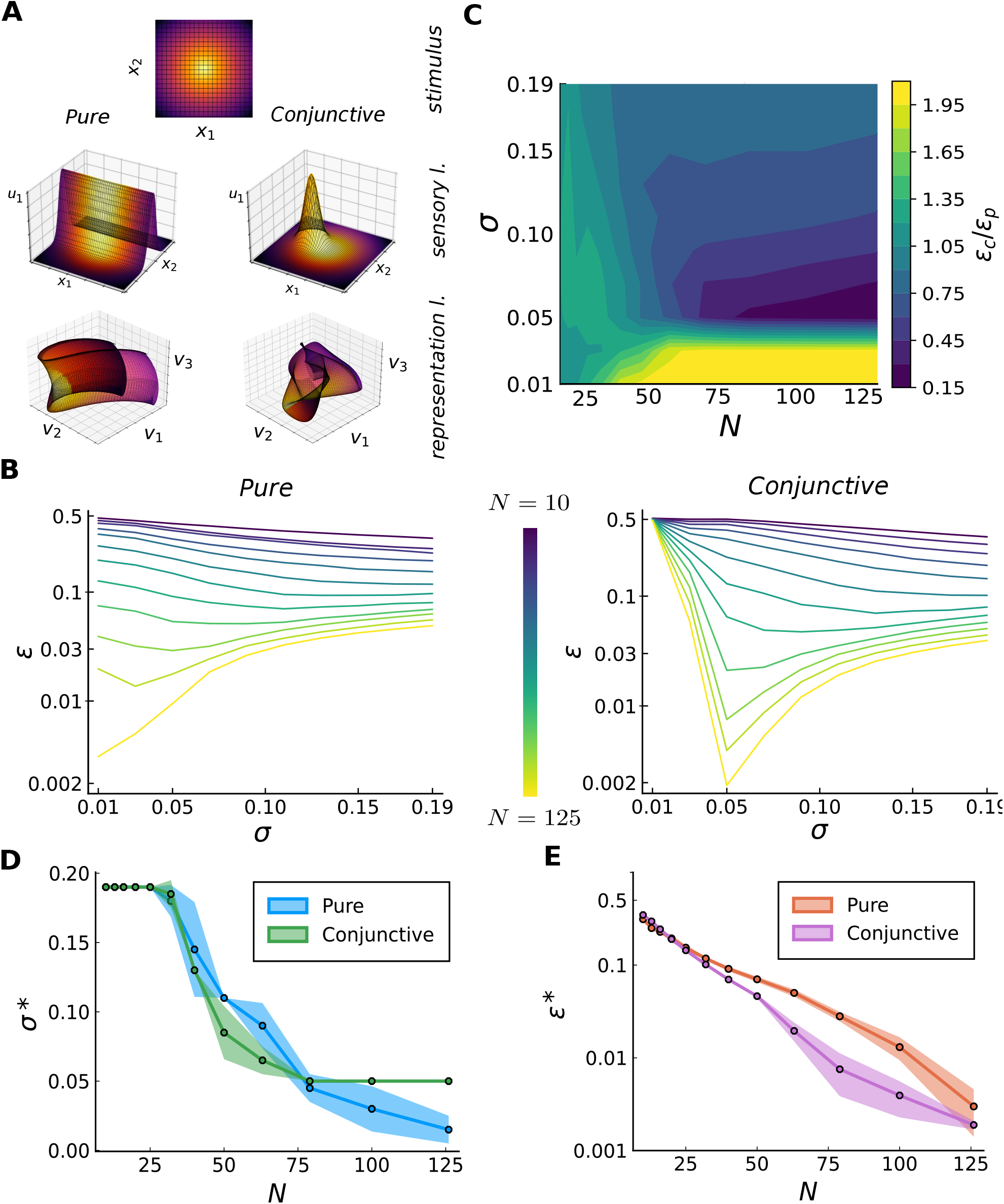
Compressed coding with multi-dimensional stimuli. We illustrate the case of a threedimensional stimulus, with *L* = 3375, *η*^2^ = 1, *R* = 1. *continue to next page* (**A**) Mapping of a multi-dimensional stimulus space into neural population activity as obtained from a two-layer coding scheme. Top: two-dimensional stimulus space; colors serve as stimulus legend for subsequent plots. Middle: mean activity (z-axis) of a sample sensory neuron, for two cases, as a function of the two stimulus coordinates (x- and y- axis). In the pure case (left), a single sensory neuron ‘folds’ the two-dimensional sheet across a direction, specified by its preferred position and dimension (here, *x*_2_). In the conjunctive case (right), a sensory neuron creates a ‘bump’ in the sheet. Bottom: joint activity of three representation neurons as a function of the stimulus. Each of these neurons randomly sum the output of sensory neurons, producing a randomly ‘folded’ sheet in the pure case (left) and a ‘crumpled’ sheet in the conjunctive case (right). (**B**) RMSE as a function of *σ* for different population sizes *N* (increasing from violet to yellow), when the first layer consists of pure (left) or conjunctive (right) cells. The optimal *σ*, which decreases with *N*, optimizes the balance between local and global errors, similarly to the one-dimensional case. In the conjunctive case, the rapid increase of the RMSE below *σ* = 0.05 is due to the sensory neurons not tiling the stimulus space, and it is independent of *N*. (**C**) Ratio of the RMSE in the two cases, *ε_c_/ε_p_*, as a function of *σ* and *N*. The yellow (violet) region indicates an outperformance of the pure (conjunctive) population. To aid visualization, the yellow region indicates all the values greater than 2. This regime of small *σ* is characterized by a better coverage of the pure population, independently of *N*. Values greater than one occur also when *N* is small, due to the prefactor of the global error being lower in the pure case. As soon as *N* is sufficiently large and *σ* is sufficiently large to allow for coverage of the stimulus space, the conjunctive case outperforms the pure case. This effect is stronger in the small-*σ* region, due to the slower scaling of the global errors in the pure case. When *σ* is large, the ratio saturates at the value given by the ratio of the local errors. (**D,E**) Optimal tuning width (**D**) and relative RMSE (**E**), for pure (blue, red) and conjunctive (green, violet) cases. The global error decreases more slowly in the pure case. For *N* ≳ 75 the optimal width in the conjunctive case saturates, due to loss of stimulus coverage, while the pure population does not suffer from this limitation. Thus, the RMSE in the conjunctive case stops decreasing exponentially and starts decreasing only linearly with *N*.

This geometric picture offers an intuitive explanation of the behavior of the MSE in the two coding schemes. (For the corresponding mathematical treatment, see Methods.) The local error is determined by how much the ‘response sheet’ is stretched out; in turn, the more the response sheet is stretched out, the more it has to fold (or crumple) to fit in the allowed range of neural activity. Folding allows for a more modest stretching of the sheet than crumpling, and as a result the pure scheme incurs a larger local error than the conjunctive scheme (see Eqs. (69) and (74)). The behavior of the global error is also different in the two coding schemes; there are two mechanisms at play, here. First, in the pure scheme, for most realizations of the random tuning curves, global errors occur primarily in a single stimulus dimension (see Methods for mathematical details); this is also apparent in Fig. 5A: the ‘folded’ structure of the response sheet induces global errors in a single stimulus dimension. By contrast, in the conjunctive scheme global errors occur in an arbitrary number of stimulus dimensions. Second, the *total* variance of the tuning curve across the stimulus space is fixed (and, in particular, set to the same value for the pure and conjunctive schemes), but the signal-to-noise ratio which governs the rate of error suppression with *N* scales differently as a function of *K*. Both mechanisms, in a regime in which *N* is large enough to suppress the contribution of global errors, enhance the probability of global error in the pure scheme as compared to the conjunctive scheme (compare Eq. (79) and Eq. (81) in Methods). Intuitively, this is because a folded sheet has a larger surface area of contact with itself than a crumpled sheet. Thus, for sufficiently large values of *N*, the conjunctive scheme is more favorable than the pure one. The corresponding crossover value of *N*, however, depends on *K*, and large values of *K* impose a stringent constraint in the conjunctive case.

We illustrate these conclusions with numerical results in the case of a three-dimensional stimulus (*K* = 3), relevant to the data analysis we present in the next section. In Fig. 5B, we illustrate the behavior of the RMSE as a function of *N* and *σ* for the pure and conjunctive coding schemes. In order to quantify the relative advantage of one scheme with respect to the other, we plot the ratio of the RMSE in the two schemes as a function of *N* and *σ* (Fig. 5C). The resulting, relatively intricate pattern, can be understood by considering different regimes. If the population size is small, the pure scheme slightly outperforms the conjunctive one (not because of a different scaling with *N*, but instead because of a difference in the prefactors that affect the probability of error in the two cases); in this regime, global errors dominate and coding is poor overall. At larger values of *N*, the contribution of local errors becomes non-negligible. If local errors dominate relative to global errors (which occurs for large *N* and sufficiently large *σ*), then the conjunctive scheme outperforms the pure one, and the ratio of the RMSEs approaches the ratio between local errors only (Eq. (75) for *K* = 3, implies 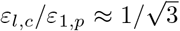). In the non-trivial regime in which local and global errors are balanced (for large N and intermediate values of *σ*), the advantage of the conjunctive scheme is further boosted. As explained above, this is due to a stronger suppression of global errors as a function of *N* in the conjunctive case. Finally, if *σ* becomes smaller than a crossover value that depends on the number of sensory neurons, the latter no longer cover the stimulus space sufficiently densely, and the conjunctive scheme breaks down; in this regime, thus, the pure scheme is favored.

As illustrated in Fig. 5B, similar to the one-dimensional case there exists in each of the two coding schemes an optimal value of the tuning curve width, *σ*, which achieves a balance between local and global errors, and it decreases with *N*. This dependence is somewhat different in the two coding schemes (Fig. 5D), and contributes to the form of the suppression of the RMSE in the two schemes (Fig. 5E). Both quantities, the optimal tuning curve width and the RMSE, decrease more rapidly as a function of *N* in the conjunctive scheme. This results from the fact that global errors are suppressed more strongly with *N* in the conjunctive case (as explained above), and therefore a smaller *σ*, yielding a lower local error, is preferable. At the same time, the requirement that sensory neurons cover the stimulus space yields a more stringent constraint on *σ* in the conjunctive scheme, yielding a bound on the extent of the regime of exponential error suppression.

### Compressed coding in monkey motor cortex

The activity of neurons in the primary motor cortex (M1) of monkey is correlated with the location and movement of the limbs. Here, we consider spatial tuning in the context of a ‘static task’ (Kettner et al., 1988). In this task, the monkey is trained to keep its hand motionless during a given delay after having placed it at one of a set of preselected positions on a three-dimensional grid labeled by the vector **x** = (*x*_1_, *x*_2_, *x*_3_). Tuning curves of hand-position selectivity can be extracted from recordings in M1 (Kettner et al., 1988; Wang et al., 2007), and it has been customary to model these as a linear projection of the hand position onto a so-called ‘preferred vector’ or ‘positional gradient’, **p**, which thus points in the direction of maximal sensitivity (Wang et al., 2007). The tuning curve of neuron i is then written as

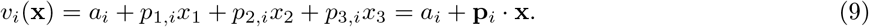

A recent study (Lalazar et al., 2016) noted, however, that a model of tuning curves that includes a form of irregularity yields an appreciably superior fit to the simple linear behavior of Eq. (9). This more elaborate model bears similarity with our model of irregular tuning curves, and this naturally led us to ask about potential coding advantages that a complex coding scheme may afford M1.

To be more specific, one can interpret the first layer in our network featured with neurons with threedimensional Gaussian tuning curves, as representing neurons in the parietal reach area (or premotor area), which are known to display spatially localized tuning properties (Andersen et al., 1985). This population of neurons projects onto a smaller population of M1 neurons which display spatially extended and irregular tuning profiles. In fitting our model to recordings from M1 neurons (Lalazar et al., 2016), we considered the arrangement of stimuli used in the experiment, namely 27 spatial locations arranged in a 3 × 3 × 3 grid fitting in a 40 cm-high cube. We then followed a previous fitting method (Lalazar et al., 2016; Arakaki et al., 2019): given the diversity of the irregular tuning curves in the population we did not aim at fitting individual tuning curves; instead, we allowed for randomly distributed synaptic weights (as in our original model) and we fitted a single parameter, the width of the tuning curves in the first layer, *σ*. The fit was aimed at reproducing specific summary statistic of the data referred to as *complexity measure* (a discrete version of the Lipschitz derivative that quantifies the degree of smoothness of a curve, see Methods and Lalazar et al. (2016)). The complexity measure varies from neuron to neuron, and we chose *σ* so as to minimize the Kolmogorov-Smirnov distance (see Eq. (103) in Methods) between the distribution implied by our model and the one extracted from the data. While our model is somewhat simpler than a model of irregular M1 tuning curves employed previously (Lalazar et al., 2016), it yields comparable fit.

With a neural response model in hand, we can evaluate the coding performance; to do so, we consider a finer, 21 × 21 × 21 grid of spatial locations as our test stimuli. We quantify the merit of a compressed code making use of irregular tuning curves by computing the MSE, 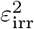, and comparing the latter with the corresponding quantity in a coding scheme with the smooth tuning curves defined in Eq. (9), 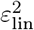. We plot our results in terms of the ‘mean fractional improvement’, Δ*ε* ≡ (*ε*_lin_ – *ε*_irr_)/*ε*_lin_. Δ*ε* is positive when irregularities favor coding, and is at most equal to unity (in the extreme case in which irregularities allow for error-free coding).

We explore the performance of the two coding schemes for different values of the parameters *N* and *σ*, first in an ideal case in which all neurons have the same noise variance (Fig. 6A). We note the existence of a crossover value of *N, N*\*. When *N < N*\*, small values of *σ* induce prohibitively frequent global errors in the compressed (irregular) coding scheme, and linear (smooth) tuning curves are more efficient. For *N > N*\*, however, irregularities are always advantageous, and the more so the smaller the value of *σ*. Because global errors are suppressed exponentially with *N, N*\* typically takes a moderate value which depends on the magnitude of the noise; the larger the noise, the larger *N*\*. Figure 6B illustrates this noise-dependent behavior of the crossover population size, for the best-fit value of *σ* (≈ 23).

**Figure 6:**
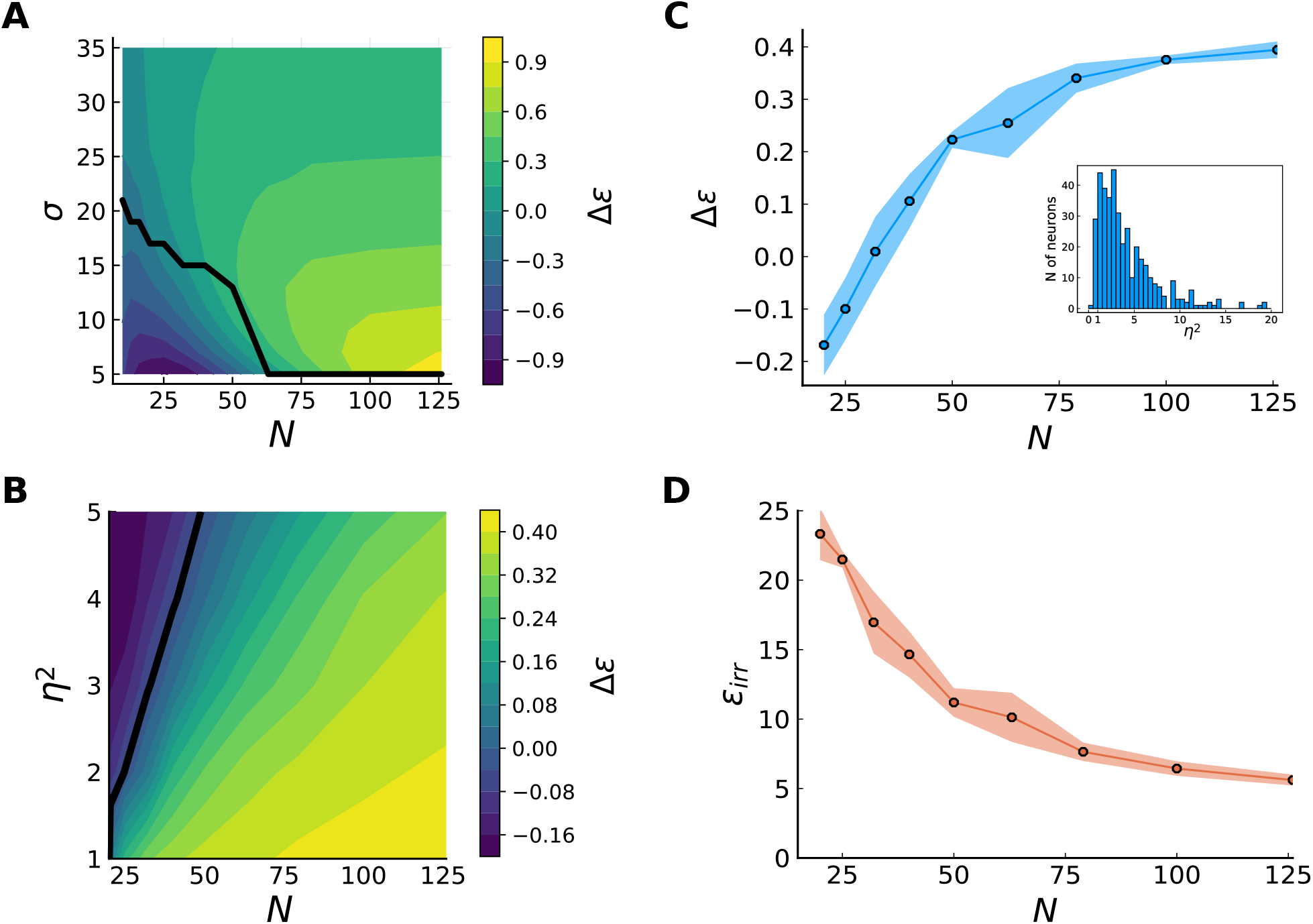
Irregular vs. linear tuning. (**A**) Mean fractional improvement for irregular tuning as compared to linear tuning, as a function of population size and tuning-curve width. The black line indicates the critical values of *N* and *σ* at which the two coding schemes perform equally well. In the region below (violet), global errors penalize the irregular case, making a smoother code more efficient. With increasing *N*, global errors become rarer while irregularities improve the local accuracy of the code (yellow region). This advantage increases at smaller values of *σ*, but so does the value of *N* required for the irregular case to be advantageous. (**B**) Mean fractional improvement in the irregular case, generated with the data-fitted model, compared to the linear one, as a function of *N* and *η*^2^. At small population sizes, irregular tuning curves produce global errors, and smoother tuning curves perform better (violet region, Δ*ε* < 0). By increasing *N*, global errors are suppressed and irregularities improve the local accuracy (yellow region, Δ*ε* > 0). The black line marks the transition values. (**C,D**) Mean fractional improvement (**C**) and RMSE (**D**) in the irregular case as a function of population size, for the noise model extracted from data. A noise variance is assigned to each neuron according to the distribution extracted from the data, showed in the inset of panel (**C**). For small *N*, linear tuning yields a better coding performance. At *N* ~ 40, the higher local accuracy compensates for global errors, and the irregular code starts to perform better, although the error is still substantial. The improvement saturates to a finite value of ~ 0.4 at a value of *N* ~ 100, when global errors are fully suppressed; the scaling of the error as a function of the population size is no longer exponential, but only hyperbolic.

Next, for a more realistic modeling of M1 neurons, we analyzed the performance of a model in which each neuron’s noise variance is extracted from data (Figs. 6C and D). For each recorded neuron, we computed the variance of the signal as the variance, across different stimuli, of the mean firing rate (left hand side of Eq. (3)). Then, we estimated the variance of the noise by averaging the trial-to-trial variability of responses to the same stimulus. These two quantities allowed us to define a signal-to-noise ratio for each neuron of the population (see Eq. (104) in Methods). As in simulations we set the variance of the signal for each neuron to a constant value, we modeled the heterogeneity in the signal-to-noise ratio as a heterogeneous noise variance; the resulting distribution is skewed, with an appreciable fraction of neurons exhibiting low signal-to-noise ratios (Fig. 6C, inset). For each value of *N*, we sampled eight different pools of *N* neurons from the population, and we averaged the corresponding mean fractional improvement, Δ*ε*. We found, again, that the relative merit of compressed coding (with irregular tuning curves) grows with the population size; interestingly, when compressed coding becomes advantageous (Δ*ε* > 0 in Fig. 6C), the error magnitude is still appreciable (Fig. 6D). This means that even though local and global errors are balanced, both contributions are substantial. Δ*ε* continues to grow with *N* until global errors are suppressed; beyond this second crossover value, *N*_local_, Δ*ε* saturates because in both coding schemes (with irregular and linear tuning curves) local errors dominate. Correspondingly, the MSE scales differently for *N* above or below *N*_local_. When *N < N*_local_ the MSE decreases exponentially with *N*, due to the suppression of global errors, while when *N > N*_local_, the suppression of the MSE is hyperbolic in *N*, reflecting the behavior of local errors only (Fig. 6D). This second crossover occurs at *N_local_* ≈ 100, a figure comparable to the number of neurons that control individual muscles in this specific task, as estimated from decoding EMG signals corresponding to individual muscles from subsets of M1 neurons (Lalazar et al., 2016).

### Dimensionality of a compressed neural code

We discussed a geometrical interpretation of a neural population code in terms of a map from a set of stimuli to a set of points in the space of (mean) population activity. With smooth tuning curves, a continuous *K*-dimensional stimulus is represented as a *K*-dimensional hypersurface embedded in the *N*-dimensional space of neural activity. This hypersurface is often referred to as a ‘neural response manifold’ (Seung & Lee, 2000; Gallego et al., 2017) (which implicitly assumes a local homeomorphism to a Euclidean space). In the previous sections, we analyzed the way in which the geometrical properties of the response manifold affect the coding performance. In this section, we relate the picture put forth by our model to recent work that quantified the dimensionality of neural activity as a way to characterize its nature and to infer strategies used by the brain to represent (sensory) information (Fusi et al., 2016; Recanatesi et al., 2020).

While a *K*-dimensional stimulus space may correspond to a *K*-dimensional neural response manifold, the latter’s complicated geometry—as in our model—may make its identification difficult. In practice, one is faced with a data set, namely a noisy sample from a population of tuning curves, and from it one would like to make statements on the geometry of the population activity. Fitting a low-dimensional manifold to a neural population data set is not a trivial task, and is the focus of a large number of studies on dimensional reduction (Cunningham & Yu, 2014). A simple approach is to consider the eigenvalue spectrum of the covariance matrix of the neural responses across the stimulus range or, equivalently, the variance carried by the the different modes in a principal component analysis (PCA). If we apply this approach to the population response in our model, for different values of *σ*, we find a spectrum that exhibits a band-pass structure, which plateaus up to a cut-off value before a sharp suppression; the cut-off value is larger for smaller values of *σ* (Fig. 7A). From this analysis one would conclude that the population activity occupies a low-dimensional subspace embedded in the N-dimensional space of neural activity, with dimensionality controlled by *σ*. As the value of *σ* falls to zero, the population responses fill an increasingly large fraction of the N available dimensions, until they fill the space entirely for *σ* → 0.

**Figure 7:**
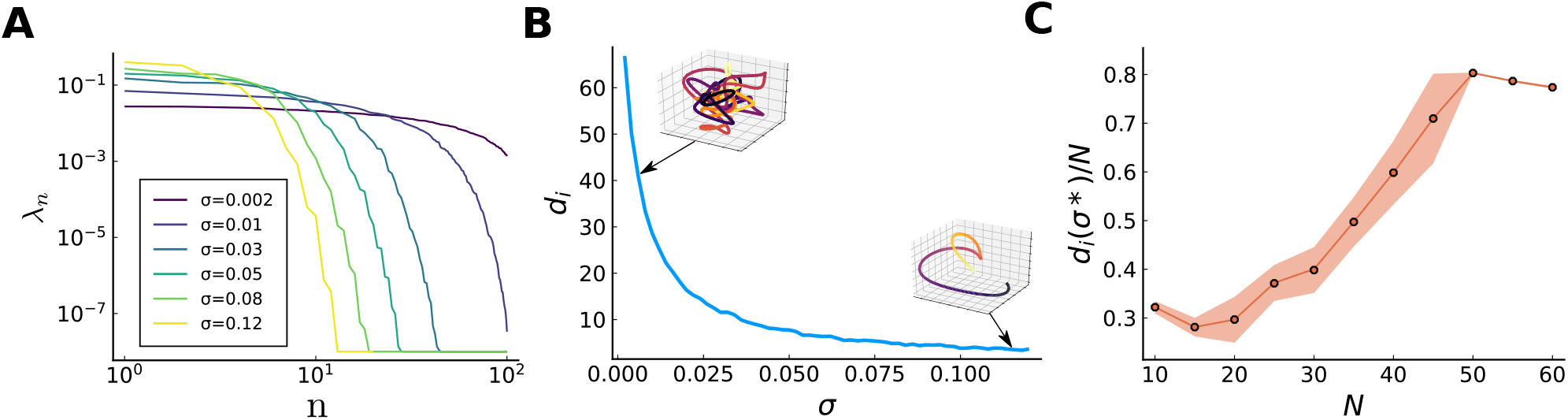
Dimensionality of the neural code. (**A**) Spectrum of the eigenvalues of the covariance matrix of neural responses in a population with *N* = 100 representation neurons, for different tuning widths (decreasing from violet to yellow). (**B**) Intrinsic dimensionality, defined as the participation ratio of the eigenvalues of the covariance matrix, in a population with *N* = 100 representation neurons, as a function of *σ*. Insets exhibit a typical response manifold in a three-dimensional space. (**C**) Ratio of the intrinsic dimensionality at the optimal value of *σ* and population size, as a function of population size, for the networks illustrated in Fig. 3,4.

A quantification of the ‘intrinsic dimensionality’ of the population activity based on this PCA analysis is offered by the participation ration, defined as 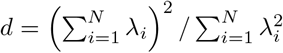, where *λ_i_* denotes the ith eigenevalue of the covariance matrix of the neural responses across the stimulus range (Gao et al., 2017). Loosely speaking, the participation ratio measures the number of eigenvalues (principal components) which are much larger than the others; for example, if *M* eigenvalues are of comparable size and much larger than any others, then *d* ≈ *M*.

In our model, while *d* is close to unity for large values of *σ*, it becomes larger for smaller values of *σ* and approaches *N* when *σ* → 0 (Fig. 7B). It is interesting to examine the behavior of this quantity in the vicinity of the optimal value of *σ*. In Fig. 7C, we display the fractional dimensionality (i.e., the participation ratio divided by the number of neurons, *d/N*) corresponding to the population activity at the optimal value of *σ* as a function of the population size, for a fixed level of noise. As expected, *d* increases with *N*: larger populations allow for more irregular tuning curves which benefit the local accuracy without generating prohibitive global errors. Quantitatively, the value of *d* hovers around *N*/2. A possible interpretation of this result is that it corresponds to the largest value beyond which a random manifold embedded in *N* dimensions comes close to intersect itself; thus, this value of d ensures that global errors do not proliferate. While a naive interpretation of the value of the participation ratio would suggest that the neural population encodes an *N*/2-dimensional stimulus, in the context of our model it results from the efficient coding of a one-dimensional stimulus. This points to the difficulty of using a simple criterion to define the dimensionality of a manifold when the latter is highly non-linear.

### Compressed coding with noisy sensory neurons

Until now, we have considered the presence of response noise only in second-layer neurons. In this case, as long as sensory neurons are tiling the stimulus space (i.e., unless there are regions in stimulus space in which sensory neurons are unresponsive), stimuli are encoded with perfect accuracy in the activity of the first layer, and the MSE inferred from activity in the second layer can be made arbitrarily small for sufficiently large *N*. If sensory neurons are also noisy, then they represent stimuli only up to some degree of precision. Furthermore, because of the (dense) projections from the first onto the second layer of neurons, independent noise in sensory neurons induces correlated noise in representation neurons. If the independent noise in sensory neurons is Gaussian with variance equal to *ξ*^2^, then the covariance of the noise in the second layer becomes **Σ** = *η*^2^**I** + *ξ*^2^**WW**^**T**^. Thus, sensory noise affects the nature of the noise in representation neurons, and it is natural to ask how this changes the population coding properties.

As we shall show, in the compression regime (*N ≪ L*) on which we focus, the kind of correlations generated by noise in the sensory layer has a negligible effect on the coding performance. The presence of sensory noise degrades coding, so a comparison of noisy and noiseless systems is not very telling. Instead, we compare population coding in the presence of the full noise covariance matrix, **Σ**, and in the presence of a diagonal covariance matrix (i.e., independent noise) with elements chosen as follows. Given the distribution of synaptic weights, the matrix **WW^T^** is sampled from a Wishart distribution with mean given by the identity matrix and fluctuations of order 1/*L* (see Methods); in the limit of *L* → ∞, the covariance matrix becomes

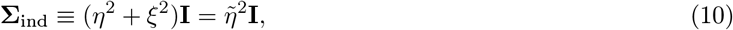

i.e., the population noise becomes independent, with single-neuron variance 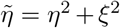. Hereafter, we compare the two cases of populations with covariance matrices **Σ** and **Σ**_ind_.

In numerical studies, we observe, first, that the MSE depends only weakly on the noise correlations, as a function of *σ*. This behavior obtains because noise correlations primarily affect local errors, not global errors. (As noise correlations reduce the noise entropy—they ‘shrink the cloud of possible noisy responses’—with respect to the independent case, one expects that correlations reduce the probability of occurrence of global errors. Numerical simulations however indicate that this effect is quantitatively negligible.)

In general, local errors can be either suppressed or enhanced by correlated noise (da Silveira & Rieke, 2021). We can show analytically that in our model, if noise correlations are due to independent noise in the sensory layer, local errors are enhanced. By computing a correction to the diagonal behavior of the covariance matrix in the limit *L* → ∞ through a perturbative expansion of the inverse covariance matrix to second order in 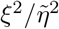 (see Methods), we obtain the local contribution to the MSE as

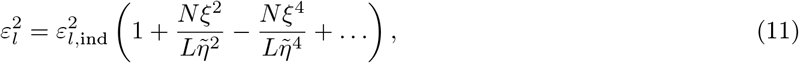

where 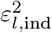 is the corresponding quantity calculated for the matrix **Σ**_ind_ rather than the full covariance matrix **Σ**. From Eq. (11), it appears that the effect of noise correlations on the MSE is deleterious, but scales proportionally to the ratio between the two population sizes, which we supposed to be small. We checked this behavior numerically (Fig. 8A), and found a good match with the analytical result. We also compared the impact of different values of *ξ*^2^, while keeping the effective noise vnoise and independent noise ofariance, 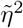, fixed (i.e., varying the relative contribution of input noise and output noise). Both Eq. (11) and Fig. 8B indicate that there exists a regime in which increasing the relative contribution of input noise, *ξ*^2^, in fact mitigates the deleterious effect of the correlated noise (this is seen in Eq. (11) as a partial cancellation of the second- and fourth-order terms).

**Figure 8:**
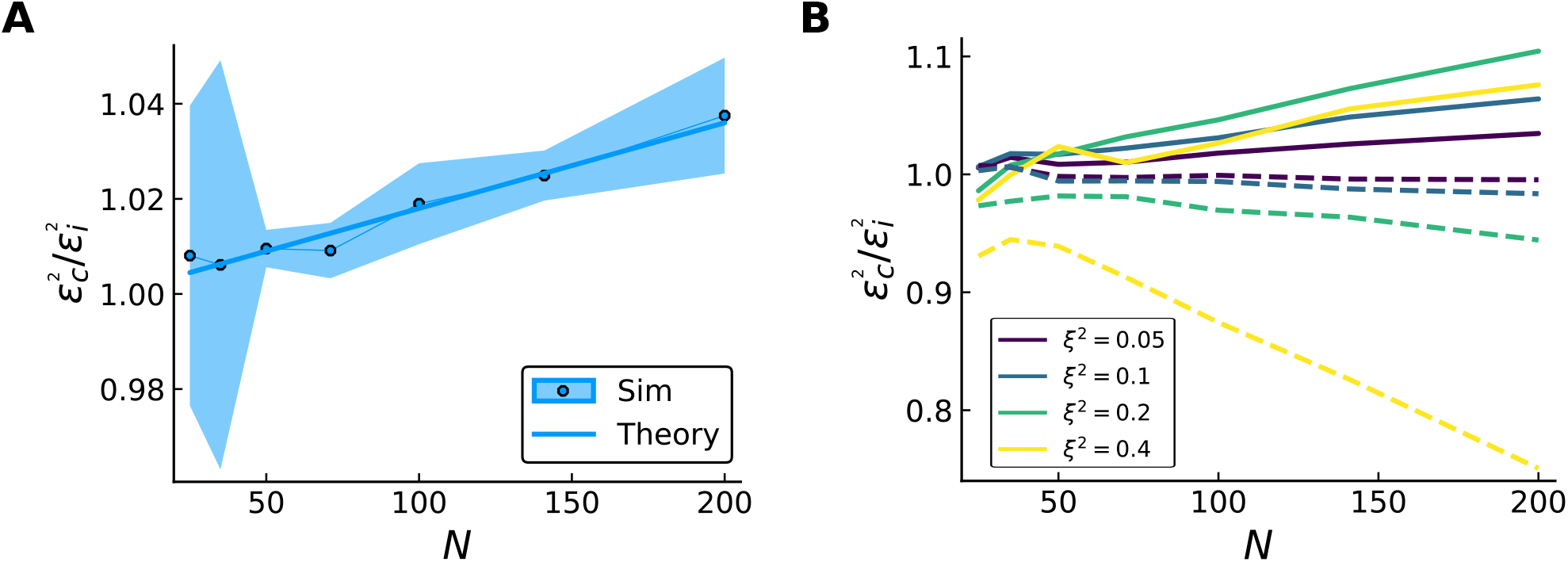
Effects of input and correlated noise on compressed coding. (**A**) Ratio of MSE in the presence of correlated noise due to input noise and independent noise of variance 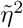, as a function of *N*, theoretical prediction (solid curve, Eq. (11)) and numerical simulations (dots) with *σ* = 0.045, 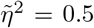 and small contribution of input noise, *ξ*^2^ = 0.05. (**B**) Ratio of MSE in the presence of correlated noise due to input noise and independent noise of variance 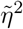 (solid curves) and ratio of MSE in the presence of correlated noise with random covariance matrix and independent noise of variance 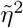 (dashed curves) Different colors denote different contributions coming from the off-diagonal terms *ξ*^2^, increasing from violet to yellow, and 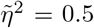. When correlations come from input noise, the ratio is positive (detrimental noise correlations). Their effect is non-linear in 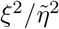, due to the competition between the first-(positive) and second-order (negative) corrections. With a random covariance matrix, correlations enhance coding precision.

Finally, we ask whether the impact of the noise correlations results specifically from the form with which sensory noise invests it. To answer this question, we examine a network with noiseless sensory neurons, but in which representation neurons exhibit correlated Gaussian noise, with a covariance matrix that has the same statistics as those of **Σ**, but in which the form of correlations is not inherited from the network structure through the synaptic matrix **W**; specifically, we consider a random covariance matrix, **Σ**_rand_ = *η*^2^**I** + *ξ*^2^**XX^T^**, where 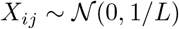. In this case, noise correlations *suppress* the MSE as compared to the independent case (with **Σ**_ind_), because the ‘cloud of possible noisy responses’ is reoriented randomly with respect to the curve of mean responses. Analytically, the analog of Eq. (11) for the case of a covariance matrix **Σ**_rand_ (instead of **Σ**) is similar, but skips the lowest-order, deleterious term:

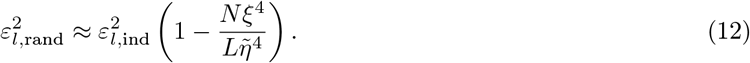

This result, as well as numerical simulations (Fig. 8B), demonstrates that generically coding is improved by random noise correlations, and that this improvement increases with *N* and also increases with the relative contribution of *ξ*^2^ with respect to *η*^2^. In sum, noise correlations in representation neurons are deleterious if they are inherited from independent noise in sensory neurons—yet, the effect is quantitatively modest.

## 3 Discussion

### Summary

We analyzed the properties of a neural population encoding a one-dimensional, continuous stimulus by means of irregular tuning curves, which emerge in a neural circuit with random synaptic weights. This model can interpolate between an irregular coding scheme, highly efficient but prone to catastrophic errors, and a smooth one, more robust in the face of noise. Optimality is achieved at an intermediate level of irregularity, which depends on the population size and on the variance of the noise. At optimality the mean error is suppressed exponentially with population size; as a result, irregular neural codes allow to compress the representation of a low-dimensional, continuous stimulus from a large, first layer of neurons to a small, second layer. We extended these results to the case of multi-dimensional, continuous stimuli, more intricate because sensory neurons can exhibit various degrees of mixed selectivity; we considered in particular a pure coding scheme, in which sensory neurons are sensitive to a single stimulus dimension, and a conjunctive coding scheme, in which sensory neurons are sensitive to all stimulus dimensions. We examined the relative advantage of one scheme with respect to the other, a question explored recently elsewhere also (Finkelstein et al., 2018; Harel & Meir, 2020), and elucidated its dependence on the number of representation neurons and on the tuning parameters. These analyses enabled us to revisit data from M1 neurons in monkey (Lalazar et al., 2016) and to discuss the benefits of an irregular code in the context of the representation of hand position. Finally, we broadened the picture of compressed coding by considering input noise, in addition to output noise, and by relating our picture to analysis of the dimensionality of population activity.

### ‘Exponentially strong’ neural population codes

Our results on the exponential scaling of the mean error with population size are similar to results obtained in the context of the representation of position by grid cells (Fiete et al., 2008; Sreenivasan & Fiete, 2011; Mathis et al., 2012; Wei et al., 2015). According to the terminology adopted in this literature, the random compressed coding presented here is an ‘exponentially strong’ population code. Grid cell-tuning is a particular instance of exponentially strong codes making use of periodicity; the model presented here offers another example, in which tuning curves are random.

The notion of an exponentially strong code predates work in computational neuroscience: Shannon introduced it in the context of communication systems and analog signals (Shannon, 1949). In his framework, a sender maps a ‘message’ (a continuously varying quantity analogous to our stimulus) into a ‘signal’ (a higher-dimensional continuous quantity analogous to the output of our representation layer) which is transmitted over a noisy channel and then decoded by a receiver. The specific illustration he provides is that of a one-dimensional message mapped into a higher-dimensional signal (Fig. 4 in Ref. Shannon (1949)), analogous to the mapping illustrated in Fig. 1C; this mapping corresponds to a curve that wraps around in a higher-dimensional space. Shannon argues that an efficient code is obtained by stretching this curve to make it as long as possible up to the point at which the winding and twisting causes the curve to pass too close to itself, thereby generating catastrophic errors.

Yet Shannon went further, and showed that such a code need not to be carefully designed. His calculation corresponds, in our framework, to the case of infinitely narrow tuning curves in the sensory layer (Fig. 2): he demonstrated that it is possible to send a discrete set of messages, with an error suppressed exponentially in the dimensionality of the signal. Our work proposes an extension of this ‘fully random’ scenario for the representation of a continuous variable based on a smooth, but irregular, mapping in a higher dimensional encoding space. By varying the width of tuning curves in sensory neurons, *σ*, one can modulate the smoothness of the mapping and trade off global errors with local errors. In this more general, ‘correlated random’ scenario, it is optimal to choose a non-vanishing value of *σ* which depends on the population size and other model parameters.

### Coding with complex tuning curves

A large body of literature has addressed the problem of coding low-dimensional stimuli in populations of neurons with simple tuning curves. The most common assumption is that of bell-shaped tuning curves; these are often chosen to model sensory coding in peripheral neurons. Various studies set in this context discussed the shape of optimal tuning curves as a function of population size and stimulus dimensionality (Zhang & Sejnowski, 1999), stimulus geometry (Montemurro & Panzeri, 2006), and the time scale on which coding operates (Bethge et al., 2002; Yaeli & Meir, 2010). More recent work analyzed the influence of a (non-uniform) prior distribution of stimuli on the optimal arrangement and shapes of tuning curves across a population of neurons; a particular prediction is that the tuning-curve width is narrower for neurons with a preferred stimulus over-represented in the prior (Wei & Stocker, 2012; Ganguli & Simoncelli, 2014; Yerxa et al., 2020). A separate direction of study focused on the effects of heterogeneity in the tuning-curve parameters on the coding performance (Wilke & Eurich, 2002; Shamir & Sompolinsky, 2006; Fiscella et al., 2015; Berry et al., 2019).

Our study falls in this line of work, but it presents two important differences: (*i*) we consider a family of irregular tuning curves (to be contrasted with simpler tuning curves, such as bell-shaped or monotonic) and (*ii*) we consider downstream neurons rather than peripheral ones. To be more specific about point (*i*), we consider tuning curves resulting from a feedforward network with random synaptic weights. The assumption of random connectivity yields a ‘benchmark model’; similar comparisons with benchmark random models have been used previously in examining information processing among layers of neural networks (Barak et al., 2013; Babadi & Sompolinsky, 2014; Litwin-Kumar et al., 2017). In our case, the irregularity of tuning curves makes the response of any single neuron highly ambiguous; the resulting code is thus distributed, and the neural population as a whole is viewed as the relevant unit of computation (Saxena & Cunningham, 2019).

Distributed codes have been argued to come with high capacity. An early example was developed in the context of face coding in the superior temporal sulcus of monkey (Abbott et al., 1996). Data analysis indicated that single-neuron sensitivity was heterogeneous and uninformative, but the number of distinguishable face stimuli grew exponentially with the population size. Our work provides an example of a random distributed code for continuous stimuli, which exhibits similar scaling properties. The main difference is that, in the case of continuous stimuli, the precise identity of the stimulus cannot be recovered in presence of noise, and what matters is the magnitude of the distance between the decoded stimulus and the true one, quantified by an appropriate metric. In other words, both the probability of occurrence of an error and its magnitude matter. The requirement of minimizing the mean squared error then yields a particular coding scheme that balances small (local) and large (global) errors.

Regarding point (*ii*), in many ‘efficient coding’ models, optimality criteria in a neural population are derived under constraints on the activity of the same population. Our results differ in that they are obtained in a downstream (‘representation’) neural population, subject to constraints on a upstream (‘sensory’) population.

### Geometry and dimensionality of population responses

In the past decade, the progress in experimental methods has allowed for the recording of neural populations on a large scale (Cunningham & Yu, 2014; Saxena & Cunningham, 2019). In an effort to interpret the way in which information is represented in population activity, various approaches have been focusing on the geometric properties of population responses to a battery of stimuli (Fusi et al., 2016; Gallego et al., 2017; Stringer et al., 2019; Kobak et al., 2019). Points in a high-dimensional space, each corresponding to the neural population response to a stimulus, are often interpreted as being located on a manifold which describes the space of possible population activity. Quantifying the geometry, and more specifically the dimensionality of this manifold, offers a characterization of neural population activity. This geometric element is eminently relevant in our work, too, where we illustrate the dependence of the coding properties of a neural population on the geometry of the representation, which in turns depends on the tuning properties of a presynaptic population (Kriegeskorte & Wei, 2021).

A specific geometrical question is that of the dimensionality of the population response in the representation layer. We showed that the spectrum of the covariance of the population activity in the representation layer, across the stimulus space, comes with a band-pass structure; by decreasing the width of tuning curves in the sensory layer, the band-pass profile acquires additional modes. Stringer et al. (2019) discussed a similar picture in analyzing recordings from a large population of visual neurons responding to a large, but discrete, set of images. In their case, the spectrum of the covariance matrix of population responses exhibits an algebraic (power-law) tail, and the authors argue that this property allows for a high-dimensional population activity while retaining smoothness of the code. Our work presents a different, and more elementary, mechanism by which a large number of modes can be accommodated by the population activity (while retaining smoothness). The non-trivial point, in our case, is that it is not beneficial for coding to be poised in the limiting case in which the number of modes is maximal but the code becomes singular (non-smooth), as, in this limit, global errors proliferate. The optimal effective dimensionality of the response manifold, as defined by the participation ratio, lies at an intermediate value at which intersections of the manifold with itself are rare and local and global errors are balanced (Fig. 7).

### Compressed sensing

We studied a network in which the information encoded in a high-dimensional activity pattern is compressed into the activity of a comparatively small number of neurons, a setting which exhibits analogies with the one of compressed sensing (Candes & Tao, 2006). Compressed Sensing is a signalprocessing approach for reconstructing *L*-dimensional signals, which are K-sparse in some basis (i.e., they can be expressed as vectors with only *K* non-vanishing elements), from *N* linear, noisy measurements, with *K* ≪ *L* and *N ≪ L* (Donoho, 2006). In our study, the low dimensionality of the stimulus, *x*, implies sparsity of the *L*-dimensional activity of the sensory layer, as long as the tuning curves in the sensory layer are not too wide.

A central result in the field of compressed sensing is that random measurements can yield near-optimal reconstructions. Furthermore, for near optimality to be achieved, the required number of measurements scales approximately linearly in *K* and only logarithmically in the dimensionality of the signal: 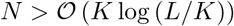 (*K*log (*L/K*)) (Candes & Tao, 2006; Baraniuk et al., 2008). In effect, in our network the representation layer operates a limited number of random measurements from the sensory layer. And we obtain an analog scaling form by inverting Eq. (5): the number of random projections, *N*, necessary to decode *L* different stimuli with negligible error scales logarithmically with the number of stimuli. We note, however, that our framework differs from that of compressed sensing as the objective is to decode the identity of the stimulus rather than a high-dimensional signal vector (in our case, the activity pattern of the sensory layer).

### Encoding vs. decoding

We focused in this study exclusively on the properties of encoding in a neural population. For this aim, throughout we assumed an ideal decoder; in principle, this is not a limitation: we show in Methods that an ideal decoder can be implemented by a simple, two-layer neural network. The first layer computes a discretized approximation of the posterior distribution over stimuli, and the second layer computes the mean of this distribution, in such a way as to minimize the MSE. Furthermore, all the operations carried out by this two-layer network—linear filtering, non-linear transfer, and normalization—are plausible biological operations (Deneve et al., 1999; Kouh & Poggio, 2008; Carandini & Heeger, 2012). The parameters involved, however, have to be chosen with the knowledge of the tuning curves and noise model.

One can ask whether biologically plausible learning rules can result in a decoder that approximates the ideal one. A closely related question has been examined by Bordelon et al. (2020), who analyzed how the generalization error in a deep neural network trained with gradient descent depends on the number of training samples and on the structure of the decomposition of a target function into a set of modes (e.g., Fourier modes). Bordelon et al. (2021) find that learning the high-frequency Fourier components of a target function requires a larger number of training samples, as compared to learning its low-frequency components. Similarly, in the context of our network one expects that learning a decoder in the case of narrow tuning curves in the sensory layer is more laborious than in the case of broad tuning curves. Noise in the training samples may also hamper learning severely in the presence of global errors. Furthermore, one can ask how our results might be modified if decoding is carried out by a decoder different from the ideal one, for example by a decoder obtained through adequately chosen learning rules. We leave these questions for future work.

## 4 Acknowledgements

We thank Anatoly Khina and Trang-Anh Nghiem for helpful discussions. Support for this research was provided by the CNRS through Unité Mixte de Recherche (UMR) 8023, the Swiss National Science Foundation Sinergia Project (CRSII5_173728), grants from the Israel Science Foundation (Grant Nos. 1319/13, 1978/13, and 1745/18), the Gatsby Charitable Foundation, and the William N. Skirball Chair in Neurophysics.

## 5 Methods

Throughout, we denote vectors by bold letters, e.g., **r** = (*r*_1_, *r*_2_,..., *r_N_*), and the *L*_2_ norm as 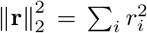. Capital bold letters, e.g., **W**, refer to matrices. We denote the derivative of a function as *f*’(*x*) = *∂f*/*∂x*.

### Network model

#### Network model for one-dimensional stimuli and constraints on its parameters

We consider a two-layer feedforward network. The first, sensory layer is made up of L neurons, each responding to a continuous scalar stimulus, *x* ∈ [0,1], according to a Gaussian tuning curve. The mean activity of neuron *j* in response to a stimulus, *x*, is given by

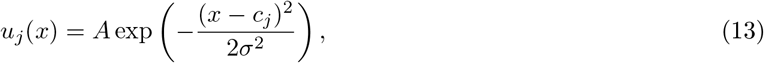

where *c_j_* is the preferred stimulus of neuron *j, σ* is the tuning-curve width, and *A* is a fixed response amplitude. The preferred stimuli are evenly spaced, *c_j_* = *j/L*. Each neuron in the first layer projects onto all *N* neurons in the second, representation layer. The transfer function is assumed to be linear, and the random synaptic weights are independent realizations of a Gaussian random variable, 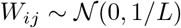; hence, the mean activity of representation neuron *i* can be written as

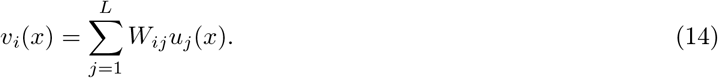

The value of the amplitude, *A*, is chosen so as to set the ‘dynamic range’ of representation neurons to a fixed value; more precisely, we choose the value of *A* so that the variance of each neuron’s response across the stimulus range is invariant under variations in the other parameter of the model, *σ*, on average over network realizations. This quantity is calculated as

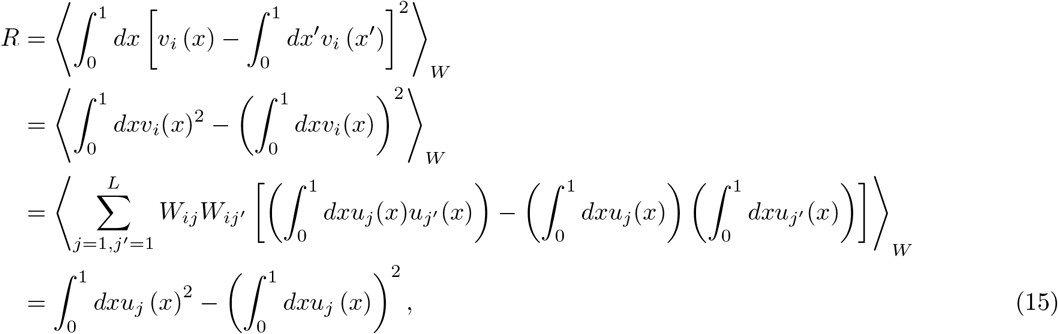

where 〈·〉_*W*_ indicates an average over the distribution of synaptic weights. Here (and below), we approximate Gaussian integrals on a bounded domain as

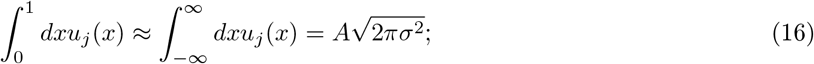

this approximation is valid when *σ* is small with respect to the stimulus range and *c_j_* is separated from the boundaries (0 and 1) by a distance that exceeds *σ*. As we will consider a large number of neurons in the sensory layer and relatively small values of *σ* (up to 1/10th of the stimulus range), errors introduced by this approximation will be negligible. By inserting Eq. (16) and a similar approximation for 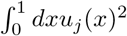 into Eq. (3), we obtain *A* as a function of *σ*, as

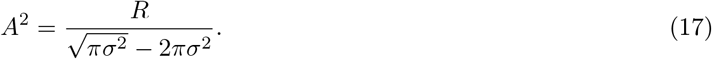

#### Tuning curves as samples from a Gaussian process

The response of each neuron in the second layer to a stimulus, *x*, is a sum of realizations of Gaussian random variables; as a result, it is also a realization of a Gaussian random variable, with mean

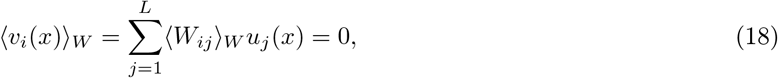

and its covariance is calculated as

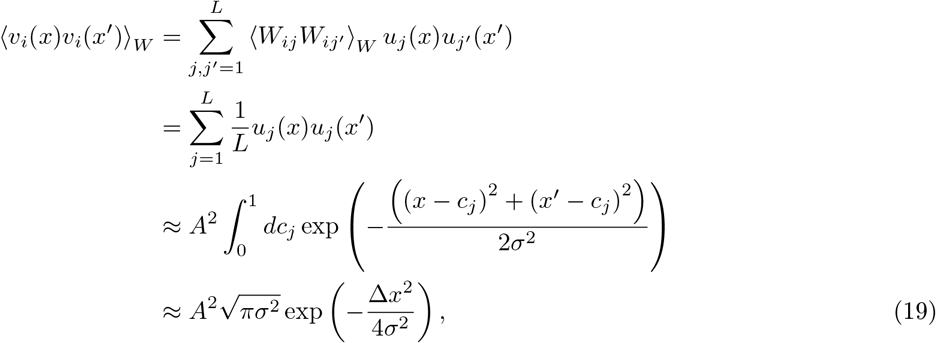

where Δ*x* = *x* – *x*’. The first approximation is obtained by replacing a sum by an integral 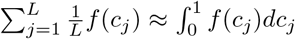 and the second approximation consists in extending the integration domain to the entire real line. The first approximation is valid if the spacing between the centers is small relatively to the width of the Gaussian, that is *Lσ* » 1, while the second is valid if the arithmetic mean of *x* and *x*’ is far from the stimulus boundaries. According to Eqs. (18) and (19) each neuron’s tuning curve can be viewed as a sample from a one-dimensional Gaussian process with vanishing mean and Gaussian kernel with standard deviation equal to 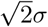 (Rasmussen, 2004).

#### Network model for multi-dimensional stimuli

We denote by *K* the stimulus dimensionality, such that the stimulus is a *K*-dimensional vector, **x** = {*x*_1_, *x*_2_,..., *x_K_*}. Analogously to the one-dimensional case, each stimulus dimension can assume values in a bounded interval, *x_k_* ∈ [0,1]. We consider the two cases of pure and conjunctive tuning for sensory neurons. In both cases, the sensory layer is made up of *L* neurons, which project onto all *N* representation neurons. Similarly to the one-dimensional case, synaptic weights are independent realizations of a Gaussian random variable, 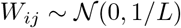.

##### Sensory neurons with pure tuning

The *L* neurons are divided in *K* sub-populations of *Q* = *L/K* neurons. Neurons in the sub-population *k* are sensitive to the single stimulus dimension *x_k_*. The mean activity of neuron *j* assigned to sub-population *k* is given by the one-dimensional Gaussian

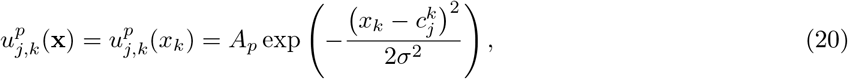

with preferred stimuli evenly spaced, 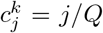 for *j* = 1,..., *Q*. The mean activity of representation neuron *i* can be written as a superposition of one-dimensional tuning curves,

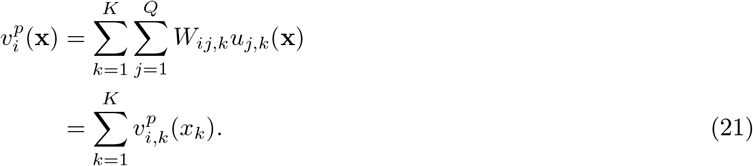

Imposing the resource constraint, Eq. (3), we obtain 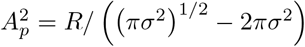.

##### Sensory neurons with conjunctive tuning

Neurons are sensitive to all stimulus dimensions. The mean activity of sensory neuron *j* is given by the multi-dimensional Gaussian function

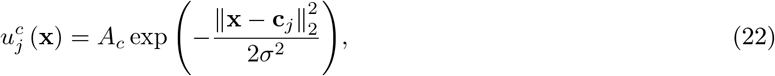

with preferred stimuli, **c**_*j*_, arranged on a *K*-dimensional square grid with mesh size *L*^-1/*K*^. The mean activity of representation neuron *i* is obtained as

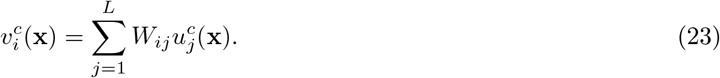

Imposing the resource constraint, Eq. (3), we obtain 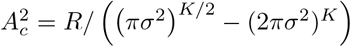.

### Population coding and optimal decoder

#### Noise Model

We assume that the response of representation neurons is corrupted by noise. The vector of responses to a given stimulus, *x*, is

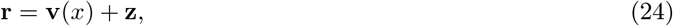

where **z** is a noise vector of independent Gaussian entries with vanishing mean and fixed variance, 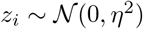. Here, **v**(*x*) = {*v*_1_(*x*), *v*_2_(*x*),..., *v_N_*(*x*)} is the vector of mean responses of second-layer neurons to a stimulus, *x* (see Eq. (2)). The probability density of a response vector, **r**, given a stimulus, *x*, is written as

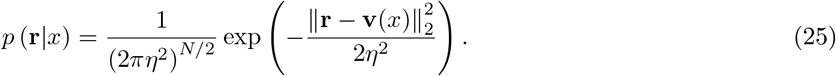

Below, we will furthermore consider an extension that takes into account a generic noise covariance matrix, **Σ**, resulting in the more general form

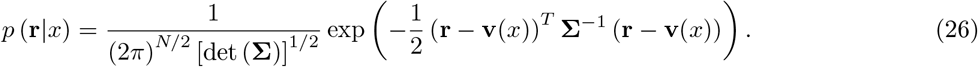

#### Loss function and ideal decoder

We quantify the coding performance of the neural population by the mean squared error (MSE) in the stimulus estimate (Dayan & Abbott, 2001), as obtained from the ideal decoder, or estimator, 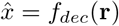, expressed as

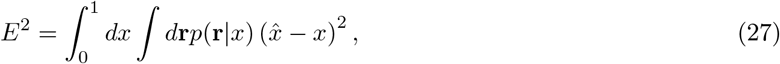

where we have assumed a uniform prior over stimuli, 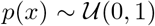. We consider the average of this quantity over network realizations, *ε*^2^ ≡ (*E*^2^)_*W*_; in some figures, we plot the square root of this quantity, the RMSE, 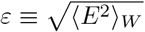.

For multi-dimensional stimuli, the ideal decoder outputs a vector estimate of the stimulus, 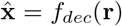. In this case, we define the MSE as the average squared norm of the difference between the stimulus and the decoder output,

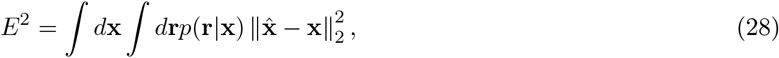

where 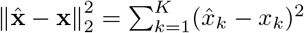.

The estimator that minimizes the MSE (Minimum-MSE or MMSE) is given by the mean of the posterior density. We can write the optimal estimator as

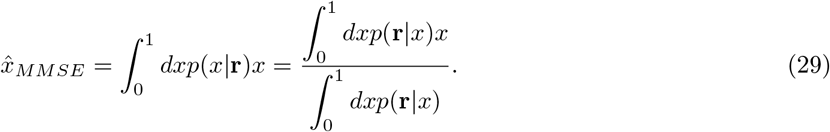

We note that a simple neural network can output the MMSE estimate. Indeed, if we approximate the integrals in Eq. (29) with a discrete sum over M values and we use Eq. (25), we obtain

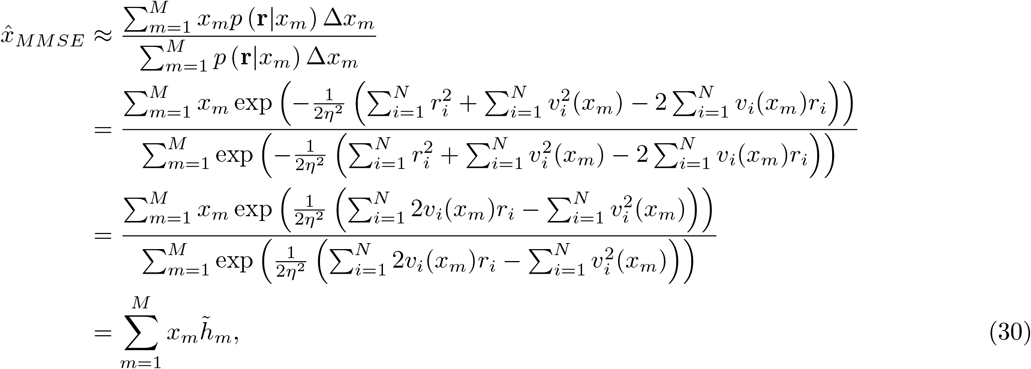

where the terms 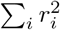 in both numerator and denominator cancel and we assumed a constant spacing, Δ*x_m_* = Δ*x*_0_. The approximate estimate specified by Eq. (30) can be by produced by a two-layer neural network: a first layer of *M* neurons, whose activities are given by

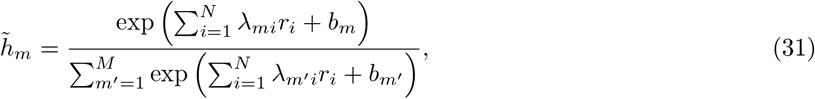

computes a normalized, discrete approximation of the posterior, 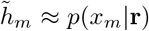, such that 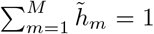. The unnormalized activity of neuron 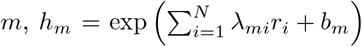, is obtained as a linear combination of the activities of the representation neurons plus a bias term, transformed through an exponential non-linearity. The ‘synaptic weight’ from the *i*th representation neuron to the mth decoder neuron is a function of the true mean response of neuron *i* to stimulus *x_m_* and of the variance of the noise, *λ_mi_* = *v_i_*(*x_m_*)/*η*^2^. Similarly, the bias term is obtained as 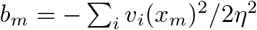. Finally, to obtain the MMSE estimate, a single output neuron weights the activity of these M neurons according to their ‘preferred stimulus’, *x_m_*.

In what follows, we will also use the maximum a posteriori (MAP) estimator, defined as

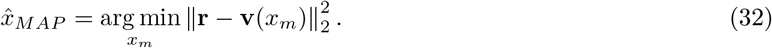

It is equal to the maximum likelihood (ML) estimator given the uniformity of the stimulus prior, and it has a simple geometric interpretation: it identifies the stimulus which corresponds to the vector of mean responses closest to the noisy population output. In numerical simulations, the MSEs calculated with the MMSE and the MAP estimators are very similar.

The MMSE estimator can be extended to the case of non-diagonal noise covariance matrix, **Σ**, by combining Eqs. (26) and (29). The decoder weights and biases are then correlated, 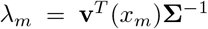 and 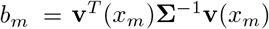, where λ_*m*_ denotes the vector with elements corresponding to the mth row of λ.

The MMSE estimator can also be extended to the case of multi-dimensional stimuli. In this case, the integrals of Eq. (29) are *K*-dimensional and the output layer is made up by *K* neurons, which compute a vector estimate of the stimulus, 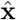.

#### Details of numerical simulations

In numerical simulations, we compute the MSE with standard Monte Carlo methods. At each step, we sample a stimulus, we generate a noisy population response and we decode it using the ideal decoder; the squared difference between the stimulus and its estimate is used to update the MSE. This process is repeated and the MSE estimate is updated until convergence, defined as the point for which the variance of the MSE estimates in the last 500 steps, after a burn-in period of 5000 steps, is less than a tolerance threshold, set to 10^-8^. We set the number of decoder neurons equal to the number of sensory neurons, *M = L*, with uniformly spaced preferred stimuli, *x_m_* = *m/M*. Unless otherwise stated, *L* = 500, *R* =1 and *η*^2^ = 0.5. The results are averaged over 8 network realizations and shaded regions corresponds to one s.d.

### Analytical derivations

In the calculations that follow, we consider the limit of *L* → ∞ and we assume *N ≪ L*.

#### Narrow tuning curves

In the limiting case with *σ* → 0, sensory neurons respond only to their preferred stimulus. Therefore, we consider the case of *L* discrete stimuli corresponding to the neurons’ preferred stimuli, *x_j_* = *j/L*. The mean activity of representation neuron i is written as *v_i_*(*x_j_*) = *ÃW_ij_*, with *Ã*^2^ = *LR*, such that 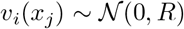. The constant of proportionality is computed with the analog of Eq. (3) for discrete stimuli, in the limit of large *L*.

The MSE in the case of narrow tuning curves, 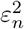, is obtained as

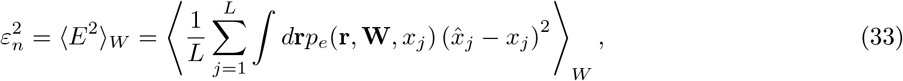

where *p_e_*(**r, W**, *x_j_*) denotes the probability, given a synaptic matrix **W** and noise, of having an incorrect estimate of *x_j_*, i.e., 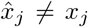. For every choice of **W** and **r**, there are *L* – 1 equiprobable realizations of the synaptic matrix which correspond to permutations of the identity of the decoded stimulus, such that 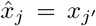 with *j*’ = *j*. Therefore, the MSE can be written as

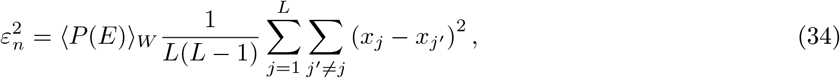

where 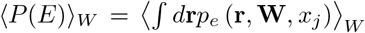 is the probability of error averaged over the noise and the synaptic weights realizations. The MSE is the product of two terms: the mean probability of error and the average squared magnitude of the error. We now compute these two terms.

##### 1 Error probability

An error occurs if there exists a *j*’ such that **r** is closer to **v**(*j*) than to **v**(*x_j’_*), where *x_j_* is the presented stimulus. We calculate the probability of error as a function of the probability of the complementary event, as

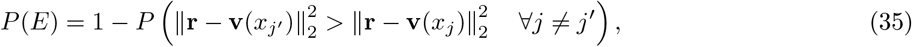

By averaging over different realizations of **W**, 〈*P*(*E*)〉_*W*_, the probabilities that an error is not committed on the possible values of *j*’ are independent; thus, we can express the probability of error, as a function of the mean responses, *v_i_*, and the noise, *z_i_*, as

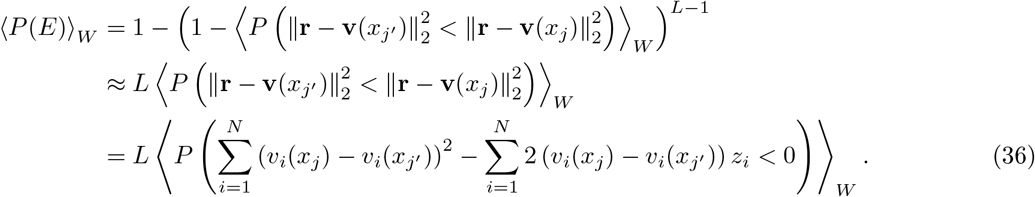

The approximation comes from the assumption that the probability of error is small, and *L* — 1 ≈ *L* is large, while the last equality is obtained from the definition of noisy responses, Eq. (24). The difference between the mean activity of the same neuron to two different stimuli is sampled according to a Gaussian distribution, 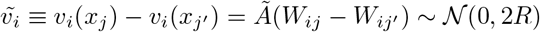. The mean probability of error is calculated as

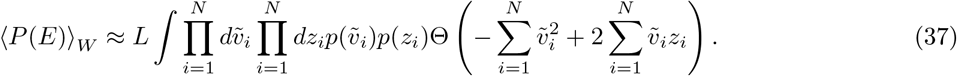

This quantity is the probability that the random variable 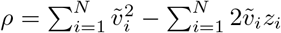, where 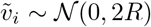 and 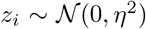, is negative. With 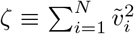, the distribution of *ρ* conditional on *ζ* is Gaussian with mean *ζ* and variance 4*ζη*^2^. Thus,

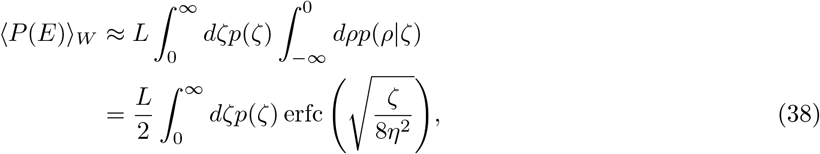

where erfc is the complementray error function and

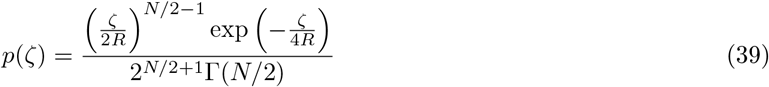

is the probability density function of a chi-squared distribution. Computing the integral, we obtain

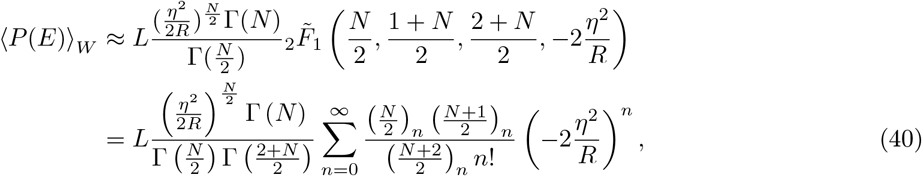

where 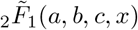 is the regularized 2F1 Hypergeometric function; we provide its definition in the last equality. The Pochammer symbol can be defined through Gamma functions, 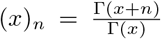. By using the identity 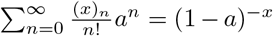 and the Stirling approximation for Gamma functions, we obtain the expression of the error probability that appears in the main text, Eq. (5):

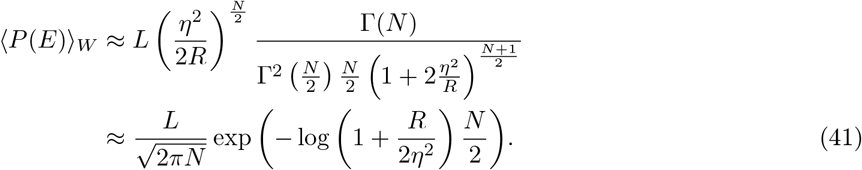

##### 1 Average squared magnitude of error

We denote by 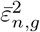 the second factor in Eq. (34), which can be written as

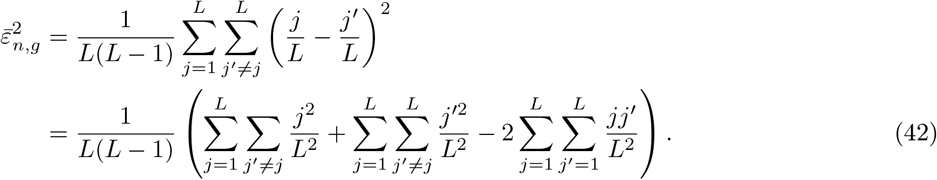

These sums can be computed through the identities for the sum of the first *n* squared numbers, 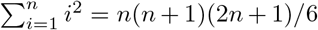, and for the sum of the first n numbers, 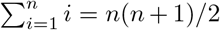. The first two sums in Eq. (42) are identical, and yield

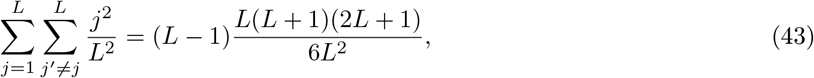

while the last term is calculated as

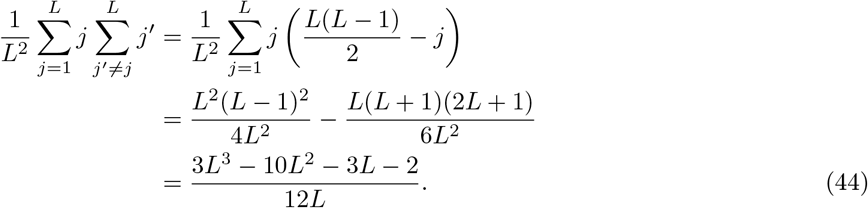

Finally, combining Eqs. (43) and (44) into Eq. (42), we obtain

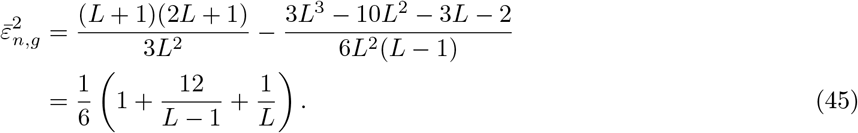

This is a term of order 1, the size of the stimulus range, plus corrections of order 1/*L*.

#### Broad tuning curves

In the case of broad tuning curves, we consider the regime of smooth response curves on the scale of the noise amplitude, such that the mean population activity can be approximated locally by a linear function of the stimulus. This regime obtains when the second-order term in the Taylor expansion is negligible with respect to the first-order one:

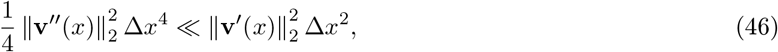

where Δ*x*^2^ ≈ *η*^2^. In order to express this condition in terms of model parameters, we impose it on average over network realizations; we note that this leads to the same result as imposing the condition on average over stimuli, but it requires a simpler calculation. From the identity 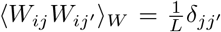, the average of the left-hand-side of Eq. (46) is obtained as

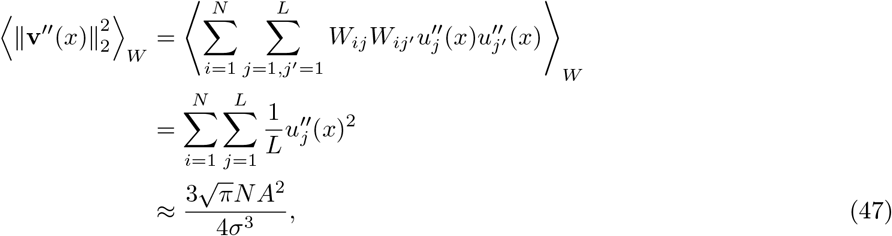

where the approximations consists in replacing the sum with an integral and in extending the integration domain to the real line. A similar calculation can be performed for the right-hand-side of Eq. (46):

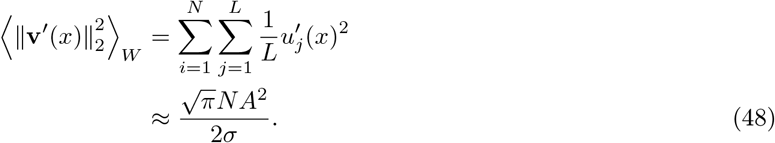

By combining Eqs. (48) and (47), and substituting Δ*x*^2^ by the variance of the noise, *η*^2^, in Eq. (46), we obtain the smoothness condition as

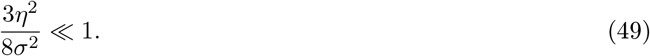

In the case of broad tuning curves the error can be of two qualitatively different types: *local* or *global* (Fig. 3A). The width of the Gaussian kernel in Eq. (19) gives a measure of the distance in the stimulus space at which population responses are correlated; we refer to a global error when the distance between the stimulus and its estimate is greater than this ‘correlation length’, *σ*. We write the MSE as

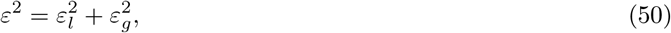

and we compute these two terms.

##### Local error

According to the ML decoder, Eq. (32), the stimulus estimate corresponds to the value *x*’ that minimizes the distance between **v**(*x*’) and **r**; if the error is local, this is obtained by projecting the noise vector onto the curve of mean population activity. By expanding the response curve around **v**(*x*), we obtain, to linear order,

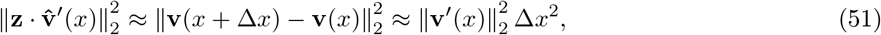

where 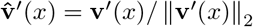. The local error can then be calculated as 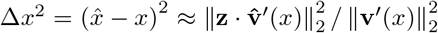. By averaging over the noise and the synaptic weights, we obtain the mean local error as

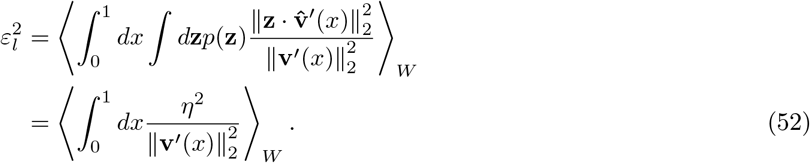

The squared norm of the derivative of the tuning curves is the realization of the random variable

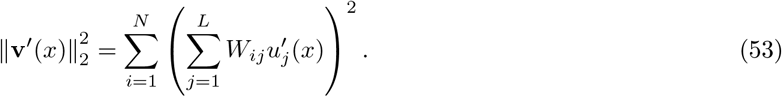

The terms of the inner sum, 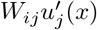, are realizations of independent Gaussian random variables with variable variance; as a result, the outer sum is also the realization of a Gaussian random variable with mean equal to the sum of the means of its terms and variance equal to the sum of the variances. The sum of the variances can be calculated as

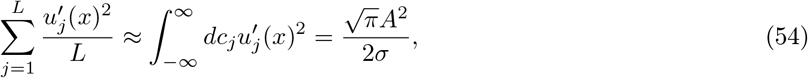

where the approximation consists in replacing the sum with an integral and in extending the integration domain to the real line. Therefore, the inner sum is distributed according to

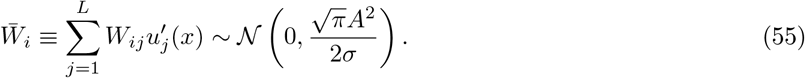

As a result, the quantity 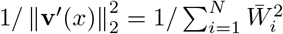 is sampled according to a scaled inverse chi-squared distribution, with mean given by

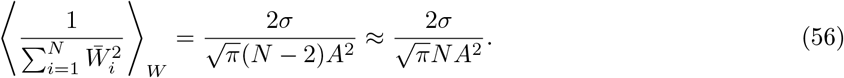

The local error is then obtained, from Eqs. (52) and (56), as

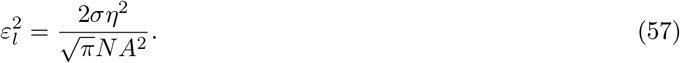

Finally, if we approximate the response amplitude for small *σ* by 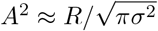, we obtain the expression that appears in the main text (first term of Eq. (6)). We note that this expression is equal to the inverse of the Fisher information averaged over network realizations; the Fisher information in case of neural responses corrupted by independent Gaussian noise is given by

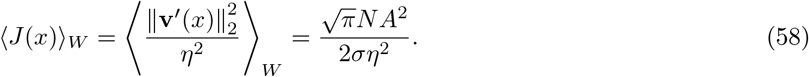

##### Global error

Here, we extend the calculation performed in the case of discrete stimuli. The analog of Eq. (34) in the case of broad tuning curves is

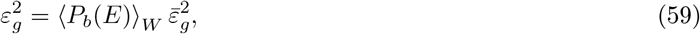

where the two factors are the probability of a global error and the average squared magnitude of a global error, respectively.

We approximate the probability of a global error by considering a division of the curve of mean population activity into *σ* ‘segments’. These segments are roughly uncorrelated and appear in random locations in the space of population activity; as a result, we can replace L by the number of segments in Eq. (5) to obtain the probability of a global error, as

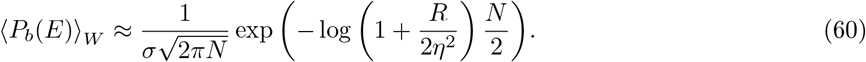

Similarly to the discrete case, when a global error occurs, the decoded stimulus is uniformly sampled from all the other stimuli belonging to incorrect segments. We illustrate the calculation for the case *x* – *σ* > 0 and *x* + *σ* < 1; similar results can be obtained for stimuli close to the boundaries of the stimulus range. In this case, the output of the decoder is distributed uniformly in the interval 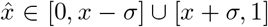. The average magnitude of global errors is therefore

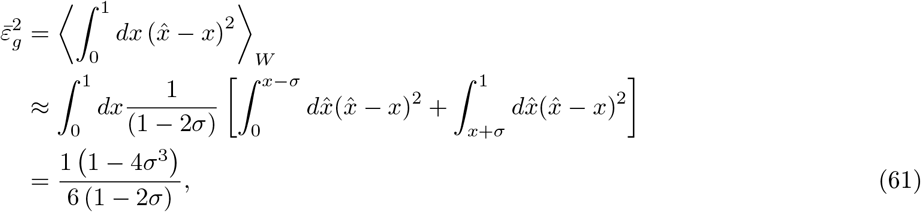

which is a term of the order 1, the size of the stimulus range, plus corrections of order *σ*. We obtain the expression for the global error which appears in the main text (second term of Eq. (6)), by combining Eqs. (59), (60) and (61).

#### Local and global errors in the case of multi-dimensional stimuli

The MSE for multi-dimensional stimuli, Eq. (28), averaged over synaptic weights realizations, is defined as the sum of the MSEs along each stimulus dimension,

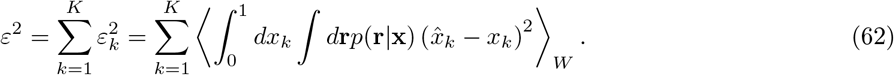

The local error along stimulus dimension *k* can be calculated, similarly to Eq. (52), as

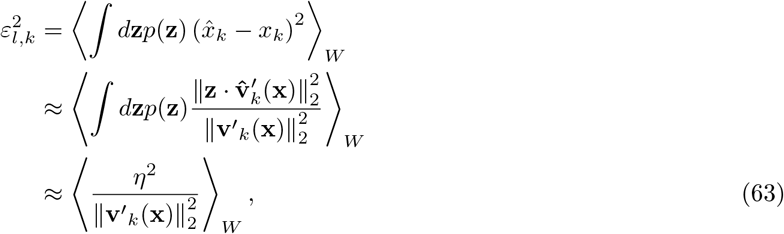

where the noise is projected onto the direction parallel to the partial derivative of the mean activity with respect to stimulus dimension *k*, **v**’_*k*_ (**x**) = *∂***v**(**x**)/*∂x_k_*

##### Local error—sensory neurons with pure tuning

The derivative of the tuning function with respect to stimulus dimension *k* is given by

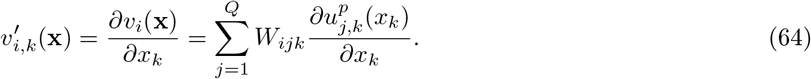

Similarly to the one dimensional case, this is a sum of realizations of independent Gaussian random variables. Dropping the superscript *p* for the sake of clarity, the sum of the variances of these terms is calculated as

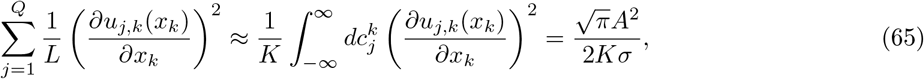

where the approximation consists in replacing the sum 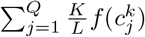 with an integral 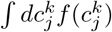, and in extending the integration domain to the real line. The sum is distibuted as

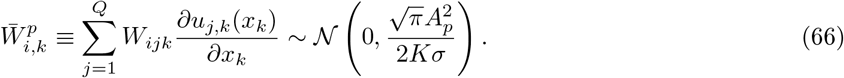

Finally, by calculating the mean of the scaled inverse chi-squared distribution,

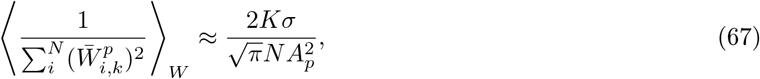

in Eq. (63), we obtain the local error along a single stimulus dimension, as

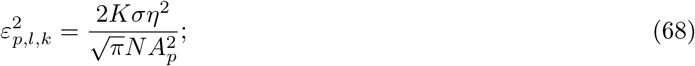

the total local error is then obtained by summing over dimensions,

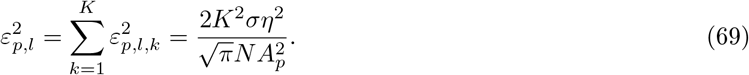

##### Local error—sensory neurons with conjunctive tuning

The derivative of the tuning function with respect to stimulus dimension *k* is given by

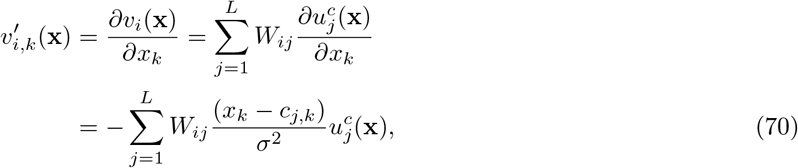

where *j* is the *k*th component of the preferred stimulus of neuron *j*, **c***_j_*. Similarly to the previous calculations, this is a sum of realizations of independent Gaussian random variables of different variances. Dropping the superscript c for the sake of clarity, the sum of the variances of these terms is calculated as

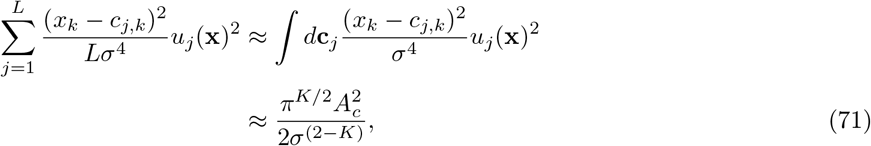

where the approximation consists in replacing the sum 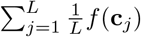 with a *K*-dimensional integral ∫*d***c**_*j*_*f*(**c**_*j*_), and in extending the integration domain. The sum is therefore distributed as

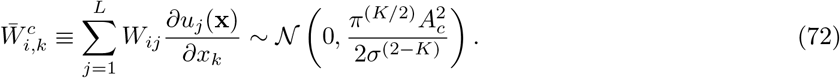

Finally, by calculating the mean of the scaled inverse chi-squared distribution,

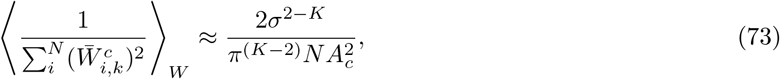

in Eq. (63), and by summing over dimensions, we obtain the total local error as

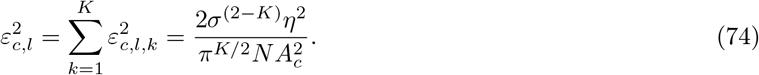

If we approximates 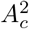 and 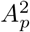 for small values of *σ*, we obtain that the ratio of the local errors in case of sensory neurons with pure and conjunctive tuning is

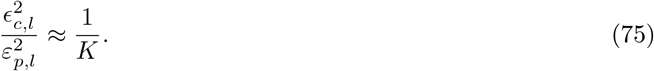

##### Global error—sensory neurons with pure tuning

In the case of sensory neurons with pure tuning, the tuning function of a representation neuron is obtained as the superposition of one-dimensional tuning curves (Eq. (21)). According to the ML decoder, Eq. (32), the decoder output can be written as

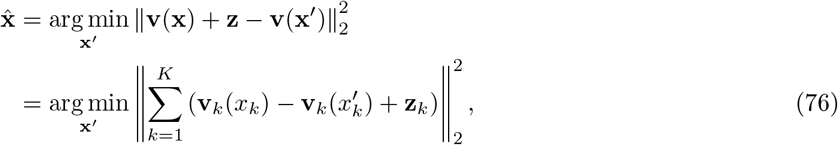

where **z**_*k*_ is the projection of the noise vector onto the direction parallel to the partial derivative of the mean activity with respect to stimulus dimension *k*. For most realizations of the random tuning curves, if *K ≪ N*, the *K* vectors summed in Eq. (76) are likely orthogonal. Thus, minimizing the squared norm of the sum is equivalent to minimizing the sum of the squared norms of each of the vectors. This, in turn, the stimulus estimate can be obtained independently for each stimulus dimension, as

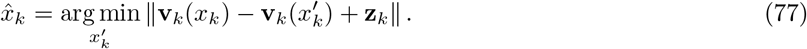

Therefore, a global error can occur in one or several stimulus dimensions; it requires that 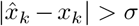 for some *k*. If the probability of a global error on more than one stimulus dimension is negligible, the total probability of a global error can be approximated as the sum of probabilities over dimensions, 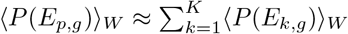. We calculated the probability of a global error in the one-dimensional case in the previous section. In order to extend the formula to this case, we have to take into account that the variance of the tuning function along one stimulus dimension is

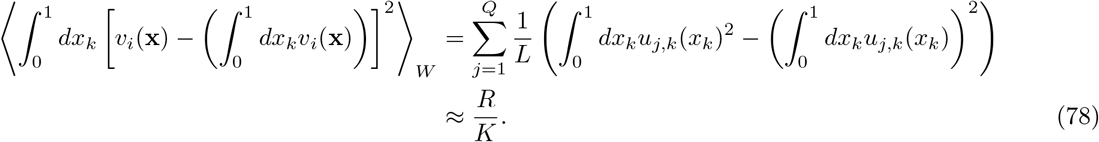

This quantity is the signal variance which governs the rate of exponential suppression of the probability of global error; replacing *R* by *R/K* in Eq. (60), multiplying by the average squared magnitude of global errors and summing over dimensions, we obtain the global error as

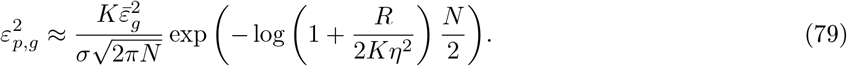

##### Global error - sensory neurons with conjunctive tuning

The correlation of the responses of neuron *i* to two stimuli, **x** and **x**’, reads

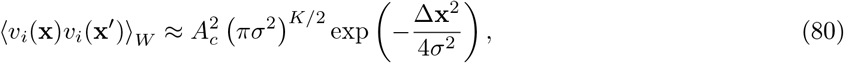

where 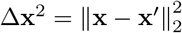; it is exponentially suppressed if 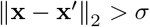. By analogy to the one-dimensional case, we divide the surface described by the population activity as a function of the stimulus, **v**(**x**) = {*v*_1_(**x**),..., *v*_N_ (**x**)}, into 1/*σ^K^* uncorrelated regions. We calculate the global error by replacing *L* with the number of uncorrelated regions in Eq. (5), obtaining

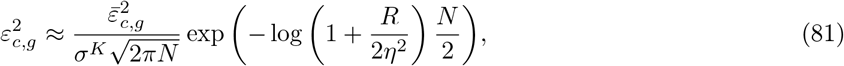

where 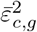 is the average squared magnitude of a global error, a term of the order of the stimulus range.

#### Influence of correlated output noise on population coding

##### Correlated output noise due to independent noise in sensory neurons

We consider the case in which the activity of sensory neurons is affected by independent Gaussian noise: 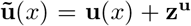, with 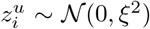. This results in a multivariate Gaussian noise in the responses of representation neurons, with covariance matrix **Σ** = *η*^2^**I** + *ξ*^2^**WW**^*T*^. The matrix **WW**^*T*^ is sampled according to a Wishart distribution, with mean **I** and variance of the matrix elements of order 1/*L* (Livan et al., 2017). We write the covariance matrix as the identity plus a perturbation, as

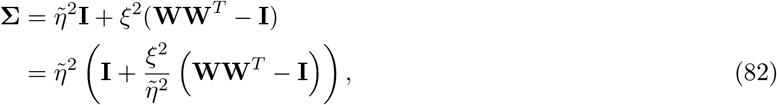

where 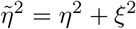. In order to quantify the effect of input noise on the coding performance, we calculate the inverse of the Fisher information (FI) as a lower bound to the MSE. The FI is written as

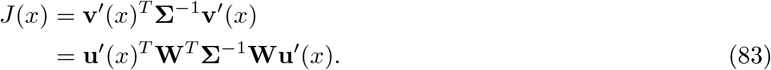

We expand the inverse of the noise covariance matrix to second order in 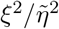, as

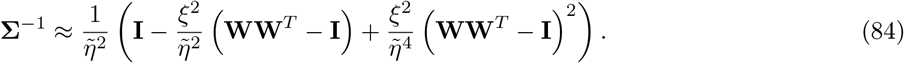

In this approximation, the FI becomes *J*(*x*) ≈ *J*_ind_(*x*) + *δJ*(*x*), with

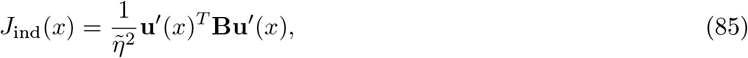

and

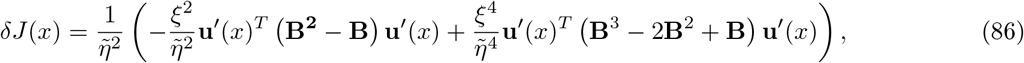

where **B** = **W**^*T*^**W**. The first term, *J*_ind_(*x*), is the FI for independent Gaussian output noise with variance 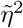; by averaging over synaptic weights realizations, we obtain the expression in Eq. (58),

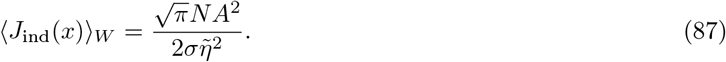

The average of the second term, *δJ*(*x*), over network realizations depends on the moments of the matrix **B**, which can be computed using Wick’s theorem: from the identity 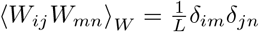, we obtain

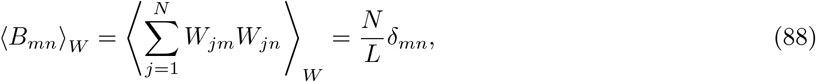

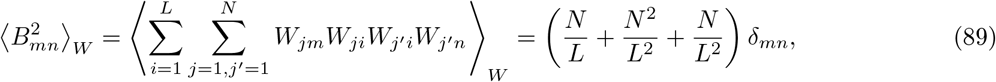

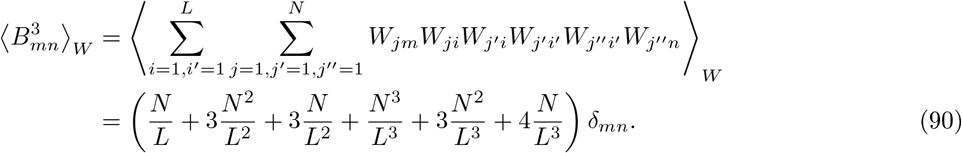

From now on, we consider the terms up to 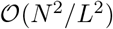; the mean of the perturbation term in the FI becomes

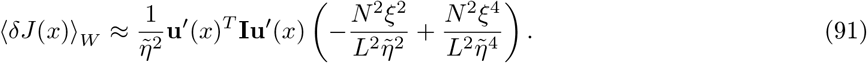

Finally, we compute the first factor by approximating the discrete sum with the integral, similarly to previous calculations, obtaining

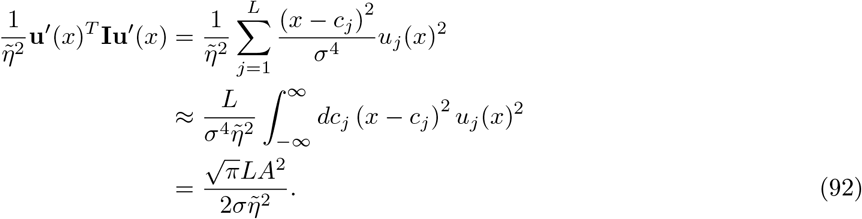

This quantity is proportional to the mean of the FI in the case of independent noise, Eq. (87), by a factor *N/L*. Combining Eqs. (87), (91) and (92), we obtain

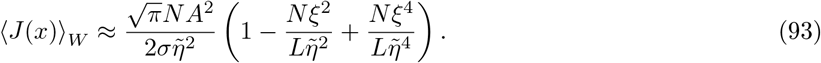

We approximate the local error as the inverse of the FI; including only corrections up to 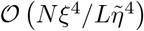, we obtain the expression that appears in the main text (Eq. (11)),

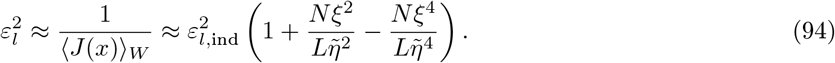

##### Correlated output noise with random covariance structure

Similar calculations can be carried out for a noise covariance matrix that obeys the same statistics as those of **WW**^*T*^, but that does not derive from the structure of synaptic weights. We consider

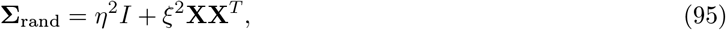

with 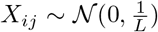, such that 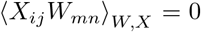 and 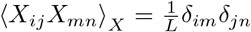. In this case, by expanding the inverse of the covariance matrix to second order in 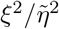 in Eq. (83), we obtain the perturbation term in the FI as

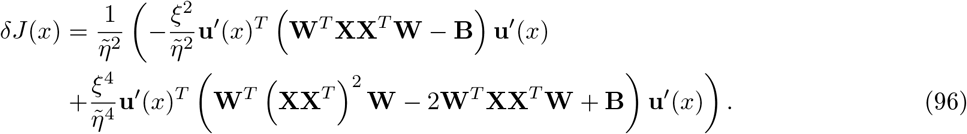

We compute the mean of these matrices over realizations of the noise covariance matrix and of the synaptic matrix using Wick’s theorem. We obtain

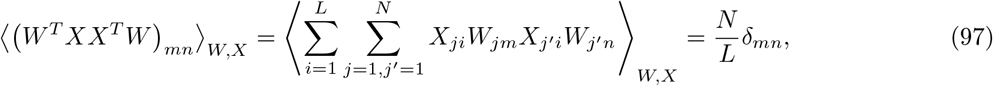

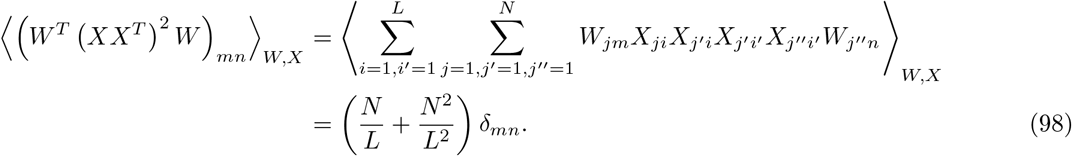

Therefore, the first order correction vanishes, and the FI is increased,

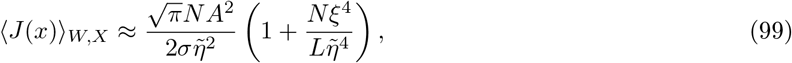

yielding a negative correction to the MSE (Eq. (12)).

### Data analysis and model fitting

#### Description of the data and summary statistics

The data consist of the responses (firing rates) of *N* ~ 500 neurons, recorded during an arm posture ‘hold’ task including 27 different positions, with 2 hand orientations each, arranged in a virtual cube of size 40×40×40 cm. The response of each neuron for each hand position is recorded in several trials (~ 10 trials per hand position). Tuning curves are computed by averaging over trials. In order to quantify the degree of irregularity of a tuning curve in a non-parametric form, the authors used a ‘complexity measure’: for neuron *i*, it is defined as the standard deviation of a discretized derivative of the mean response:

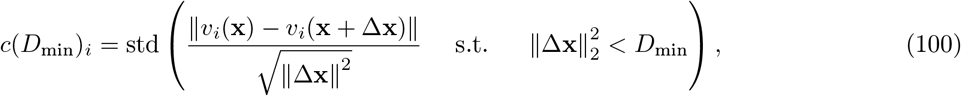

where *v_i_*(**x**) is the mean response, *D_min_* is the distance between two neighboring hand positions, and in the experiment is equal to 35. Lalazar et al. (2016) evaluated also another summary statistics, the distribution of *R*^2^ values resulting from a fit of the tuning curves with a linear model (see Eq. (9), originally proposed by Kettner et al. (1988)):

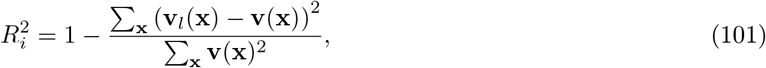

where **v**_*l*_(**x**) is the response predicted by a linear regression of the data, and the sum is over hand positions used in the experiment. The distribution of these two quantities across neurons is a measure of the irregularity of the neural population response; if the population were perfectly described by a linear model, the *R*^2^-distribution would be a constant for all neurons and equal to 1, while the complexity measure would exhibit low values.

#### Model fitting and comparison between irregular and linear tuning curves

We consider neurons responding with at least 5 spikes/s at more than two target positions and we compute their tuning curves by averaging the firing rates over trials. Then, we shift and normalize the tuning curves to cancel their means and set their variances across hand positions to unity. We use a version of our shallow network model to produce three-dimensional mean-response profiles. The sensory layer is made up of *L* = 100^3^ neurons; the preferred stimuli (here, hand positions) are arranged so as to cover a space of 100×100×100 cm, in such a way that hand positions used in the experiment are placed far from the boundaries of the stimulus space. To limit computation load, we choose **W** as a sparse random matrix, with sparsity equal to 0.1, with Gaussian-distributed elements, similarly to the model of Lalazar et al. (2016). The sparsity of the matrix does not affect our results, as long a proper normalization of the synaptic weights is taken into account and the representation neurons receive a sufficient number of inputs from the sensory layer, i.e., as long as the matrix is not too sparse and the tuning width is not too narrow. The tuning curves are normalized to have zero mean and unit variance across hand positions. With respect to the model of Lalazar et al. (2016), there are two main differences: in their case the random weights were distributed according to a uniform distribution, and a rectifying non-linear function was applied to the random sum of the activity of first-layer neurons to enforce a positive activity of the representation neurons. Their model thus had two tunable parameters: the tuning width of first-layer neurons, *σ*, and the the threshold of the non-linear transfer function in the second layer. The only tunable parameter in our model is *σ*.

In order to fit our model, we generate neural responses of a number of representation neurons equal to the number of recorded neurons, using the same set of hand positions to as used in the experiment. We then computed the distribution of the complexity measure for different values of *σ*; we denote by *σ_f_* the tuningcurve width which minimizes the Kolmogorov-Smirnov (KS) distance between the distribution produced by the model and that extracted from the data (Fig. S1A). The KS distance is a measure of discrepancy between two probability distributions. We denote by *F*_data/model_(*c*) the empirical cumulative distribution function of the complexity measure across data/model, that is, the empirical probability of finding a neuron with complexity measure less than *c*,

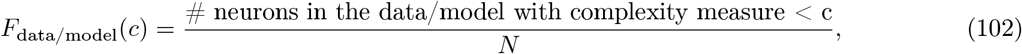

where *N* is the total number of neurons. The KS distance is defined as the maximum absolute difference between the *F*_data_ and *F*_model_:

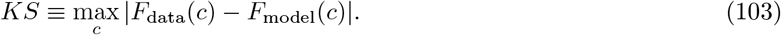

Figure S1C compares the distribution of the complexity measure across neurons for our model with *σ* = *σ_f_* with the one found in data and the one calculated for a population with linear tuning curves. For the sake of completeness, we also computed the KS distance between the distributions of *R*^2^ corresponding to model and data (Fig. S1A, red line). We mention that the model of Lalazar et al. (2016) with two tunable parameters did not reproduce the distributions of complexity measure and of *R*^2^, and only the complexity measure was taken into account in the fitting procedure. A better fit can be obtained in a heterogeneous model, at the cost of tracking many more parameters (two per neuron): see Arakaki et al. (2019) for a more detailed discussion of the fitting procedure in such a model.

We also extract a noise model from the data, as follows. We define the variance of the mean response of neuron *i* across hand positions as the variance of the average responses across hand positions, 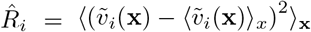, where 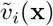 is the unnormalized tuning curve. Similarly, we average the trial-to-trial variability across different stimuli to obtain the variance of the noise, 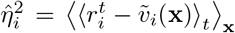, where *r*^*t*^ is the response at trial *t*. In the model, we set the variance of the signal to unity and we rescaled the noise variance correspondingly, as

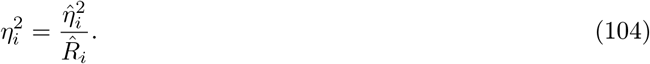

In principle, the noise may depend on the stimulus. To control for this effect, we preprocess the data with a variance stabilizing transformation. We substitute *r_i_*(**x**) by 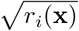, (SRJ & Everitt, 1999)), and we recalculated the variance of the noise accordingly. In this way, if the noise were proportional to the mean, one would obtain a constant estimate of the variance of the responses for different hand positions. The distribution of noise variances across neurons calculated in this way does not differ substantially from the one obtained without this data transformation.

For numerical simulations (Fig. 6), the tuning curves are computed at a finer scale than in the data (cubic grid of 21×21×21 points instead of 3×3×3). We illustrate three examples of tuning curves obtained with *σ* = *σ_f_*, measured at these hand positions in Fig. S1D-F, together with the prediction obtained from a linear regression (Eq. (9)). We note that there are some neurons which are well described by the linear model while others are not compatible with it. We generated the tuning curves for a number of neurons equal to the number of neurons analyzed in the fitting procedure (*N*_tot_ = 400). Results for a given population size, *N*, are obtained by averaging over 8 different pools of size *N* sampled with replacement from *N*_tot_. In Fig. 6A-C, we compare the MSE as obtained in a population in which neurons respond according to the irregular tuning curves generated by our model and a population in which the tuning curves are linear, Eq. (9). The latter are generated according to Eq. (9), by sampling the preferred directions, **p**_*i*_, uniformly on the unit sphere; the tuning curves are shifted and normalized to have zero mean and unit variance across hand positions. The comparison is quantified through the mean fractional improvement, defined as

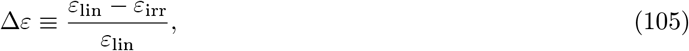

where *ε*_lin_/_irr_ is the RMSE as obtained in the population with linear/irregular tuning curves.

### Resources availability

The data we analyze were reported and discussed by Lalazar et al. (2016). Data are publicly available at https://osf.io/u57df/. Numerical simulations and data analyses were carried out with custom codes written in Julia (Bezanson et al., 2017), with the DrWatson project assistant (Datseris et al., 2020). The custom code will be made publicly available upon manuscript acceptance

**Fig. S1:**
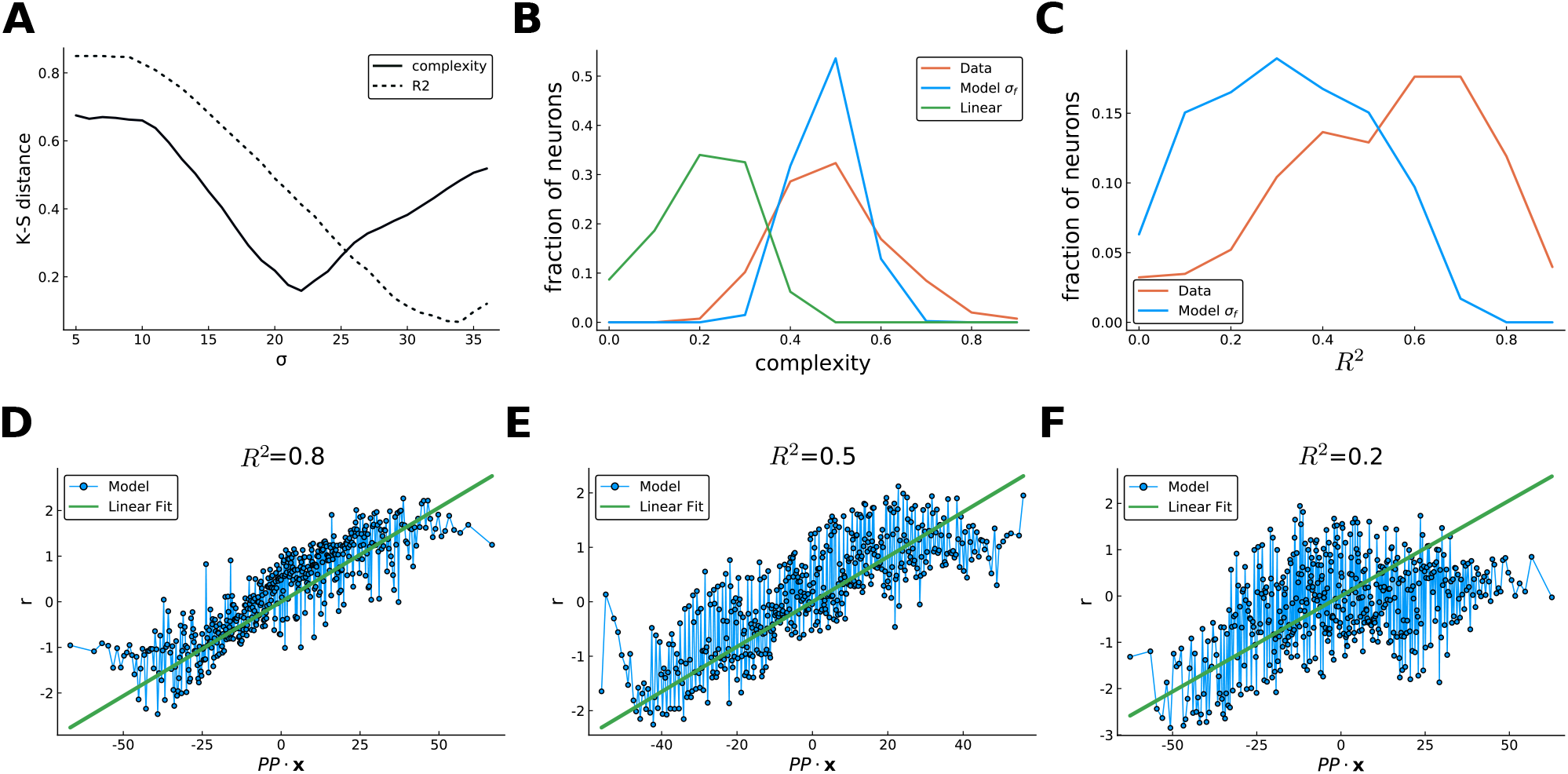
Model fitting and tuning curves. (**A**) Kolmogorov-Smirnov distances between the distributions of complexity measure (solid line) and *R*^2^ of fitting across neurons (dashed line) for data and model, for different values of *σ*: *σ_f_* is chosen to be the value at which the minimum of the distance between complexity distributions is attained, *σ_f_* ~ 22. (**B**) Normalized-histograms of the distribution of complexity measure (arbitrary units) across the neurons in the data (red), with irregular tuning with *σ* = *σ_f_* (blue) and a linear tuning curves (green). The irregular model captures the bulk of the distribution for the data better than a linear model. Nevertheless, the data present a broader distribution across the population. (**C**) Normalized-histograms of the distribution of the *R*^2^ of linear fits across neurons of the data and irregular tuning curves with *σ* = *σ_f_* (red). (**D-F**) Three examples of irregular tuning curves with *σ* = *σ_f_*, showing a broad range of behaviors with respect to a linear fit. The tuning curves are plotted as a function of the projection of the hand position onto a preferred direction, obtained by the fit with Eq.(9) (green line). Some neurons are well described by the parametric function (**D**), while others show consistent deviations (**E**); in a subset of neurons, a linear fit fails altogether (**F**)

